# High-throughput quantification of camelid cytokine mRNA expression in PBMCs by microfluidic qPCR technology

**DOI:** 10.1101/2023.01.15.524100

**Authors:** Jordi Rodon, Nigeer Te, Maria Ballester, Joaquim Segalés, Júlia Vergara-Alert, Albert Bensaid

**Affiliations:** Unitat mixta d’investigació IRTA-UAB en Sanitat Animal, Centre de Recerca en Sanitat Animal (CReSA), Campus de la Universitat Autònoma de Barcelona (UAB), 08193 Bellaterra, Catalonia, Spain; IRTA, Programa de Sanitat Animal, Centre de Recerca en Sanitat Animal (CReSA), Campus de la Universitat Autònoma de Barcelona (UAB), 08193 Bellaterra, Catalonia, Spain; Animal Breeding and Genetics Program, Institute for Research and Technology in Food and Agriculture (IRTA), Caldes de Montbui, Spain; Departament de Sanitat i Anatomia Animals, Facultat de Veterinaria, Universitat Autònoma de Barcelona (UAB), Campus de la UAB, 08193 Bellaterra, Catalonia, Spain

**Keywords:** Camelid species, cytokines, immune responses, RT-qPCR, Fluidigm Biomark, PHA, PMA-ionomycin, PolyI:C, dromedary, camel, llama, alpaca

## Abstract

Camelids are economically and socially important in several parts of the world and might carry pathogens with epizootic or zoonotic potential. However, biological research in these species is limited due to lack of reagents. Here, we developed RT-qPCR assays to quantify a panel of camelid innate and adaptive immune response genes, which can be monitored in a single run. Validation of the assays was performed with PHA, PMA-ionomycin, and Poly I:C-stimulated PBMCs from alpaca, dromedary camel and llama, including normalization by multiple reference genes. Further, comparative gene expression analyses for the different camelid species were performed by a unique microfluidic qPCR assay. Compared to unstimulated samples, PHA and PMA-ionomycin stimulation elicited robust Th1 and Th2 responses in PBMCs from camelid species. Additional activation of type I and type III IFN signalling pathways was described exclusively in PHA-stimulated dromedary lymphocytes, in contrast to those from alpaca and llama. We also found that PolyI:C stimulation induced robust antiviral response genes in alpaca PBMCs. The proposed methodology should be useful for the measurement of immune responses to infection or vaccination in camelid species.

## INTRODUCTION

Camelid species are livestock of great economic, sanitary, health and environmental importance in the north of Africa, central Asia, the Middle East, and South America. Number of these animals is expected to grow since the camelid industry is in transition from nomadism to intensive production (FAOSTAT, 2021). In Europe, camelids are used for fine wool production but also kept as pets, guardians of other livestock, or used for recreational or leisure purposes (Halsby et al., 2017). However, these species are susceptible to several viruses, bacteria and protozoan parasites affecting meat and milk production. As an example, Camelpox and Peste des Petits Ruminants viruses are causing recurrent epizootic outbreaks in Africa and Middle East (Khalafalla, 2017). Furthermore, there is limited information available about the role of camelids in the epidemiology of zoonotic diseases. In recent years, the most studied pathogens causing zoonotic diseases included the Middle East respiratory syndrome coronavirus (MERS-CoV), Rift Valley fever virus (RVFV), *Brucella* sp., and *Echinococcus granulosus* (Zhu et al., 2019). Many other less studied viruses known to be carried and transmitted by camelids, such as hepatitis E virus (HEV) or Crimean-Congo haemorrhagic fever virus (CCHFV), are of serious human health concern as well. Camelids are also hosts of many other fastidious bacterial diseases including tuberculosis, gastrointestinal illnesses caused by verotoxin-producing *E. coli*, campylobacteriosis, listeriosis and salmonellosis, among others, as well as protozoan parasites (*Cryptosporidium* spp., *Sarcoptes, Giardia duodenalis*, etc.) of veterinary and human health concern (Khalafalla, 2017).

Understanding disease pathogenesis and identifying protective immune responses are prerequisites for the rational development of new anti-microbial drugs and vaccines. In addition, comparison of immune responses between humans and domestic or wildlife species would shed insights to delineate host factors involved in disease outcome. Upon pathogen infection, the host immune system is regulated by complex mechanisms in which cytokines play a pivotal role in determining the intensity and duration of the immune response (Dinarello, 2007; Svanborg et al., 1999). In some domestic species, such as pig, goat, cattle, and sheep, the quantification of cytokines, either at the protein or mRNA level, has become a widely used method to monitor immune responses upon pathogen infection (Smeed et al., 2007; van Reeth & Nauwynck, 2000). However, cytokine detection in camelids has been hampered by the lack of specific reagents.

To date, there are few reliable commercial ELISA kits available to study immune responses in camelids at the protein level. Nonetheless, camelid interferon (IFN)-α, and some Th1 and Th2 cytokines, and pro-inflammatory cytokine cDNAs have been cloned and sequenced (Nagarajan et al., 2012; Odbileg et al., 2006; Odbileg, Konnai, et al., 2005; Odbileg, Lee, et al., 2004, 2005; Premraj et al., 2020). Sets of primers have been derived from these sequences (Odbileg, Konnai, et al., 2004) to quantify cytokine mRNAs in peripheral blood mononuclear cells (PBMCs) of *Camelus bactrianus* upon vaccination with *Brucella abortus* strain 19 by reverse transcription quantitative polymerase chain reaction (RT-qPCR) assays (Odbileg et al., 2008). Although these previous works provided tools to quantify a few camelid cytokine mRNAs, primer assays only allow for a cursory study of immune response pathways and were not optimized to function in high-throughput qPCR platforms. We previously took advantage of well-annotated camelid draft genomes (Wu et al., 2014) to design a comprehensive set of primers from genes encompassing several innate immune response pathways (Te et al., 2021, 2022), and demonstrated their usefulness in respiratory tract samples of alpacas. Here, we extended this panel of primers to characterize the expression of innate and adaptive immune response genes in phytohemagglutinin (PHA), phorbol 12-myristate 13-acetate (PMA)-ionomycin and/or PolyI:C-stimulated PBMCs from three different camelid species (dromedaries, llama, and alpacas). We optimized gene expression analyses in the highly sensitive and cost-effective Fluidigm Biomark microfluidic qPCR system. A full validation and standardization of these assays is provided together with an interspecies comparison, characterizing camelid cytokine expression with non-specific PBMC stimuli widely used in immunological research.

## MATERIALS AND METHODS

The present work was performed using the nomenclature and following the validation protocols proposed by the Minimum Information for publication of quantitative real-time PCR experiments (MIQE) guidelines (Bustin et al., 2009).

### Animal welfare, ethics, and experimental design

Experiments with animals were performed at animal facilities of IRTA Alcarràs farm (Lleida, Catalonia, Spain) or at the Biosafety Level-3 (BSL-3) facilities of IRTA-CReSA (Barcelona, Catalonia, Spain), and were approved by the Ethical and Animal Welfare Committee of IRTA (CEEA-IRTA) and by the Ethical Commission of Animal Experimentation of the Autonomous Government of Catalonia (approval No. FUE-2017-00561265 and FUE-2018-00884575).

Two llamas (L1, L2) and five alpacas (A1-5) were purchased from Belgium and The Netherlands, respectively, housed at IRTA-CReSA animal facilities and used for routine blood collection. L1 and L2 were used in a previously published study (Rodon et al., 2019), and blood was collected prior experimental infection with MERS-CoV. One healthy dromedary camel (D1) from a private zoo (Alicante, Valencian Community, Spain) was also bled once for routine checking purposes, and extra blood samples were taken to perform this work.

### Blood collection

Whole blood samples (40 to 50 mL) from each animal were collected from the jugular vein using EDTA BD Vacutainer® tubes (Beckton Dickinson, New Jersey, USA), following animal welfare protocols.

### PBMC isolation

Prior PBMCs isolation, whole blood was diluted 1:1 with phosphate-buffered saline (PBS). PBMCs were harvested from blood by density-gradient centrifugation with Histopaque®-1083 (Sigma-Aldrich, St. Louis, MO, USA), according to the manufacturer’s instructions. PBMCs were cultured in RPMI-1640 media supplemented with antibiotics (100 U/mL penicillin, 0.1 mg/mL streptomycin) and glutamine (2 mmol/L) purchased from Life Technologies (Waltham, USA), β2-mercaptoethanol (5×10^−5^ M; Sigma-Aldrich, MO, USA), and 10% heat inactivated foetal calf serum (FCS; EuroClone, Pero, Italy). Cell viability was assessed by the Trypan blue staining exclusion method.

### Cell stimulation assays

PBMCs from A1-2, D1, and L1-2 were seeded on 24-well plates at 5·10^6^ cells/mL, and cultured in duplicates in medium alone (control condition), or stimulated with 10 µg/mL of PHA (Sigma-Aldrich, St. Louis, MO, USA), or with a combination of 10 ng/mL PMA (Sigma-Aldrich, St. Louis, MO, USA) and 1 µg/mL ionomycin calcium salt (Sigma-Aldrich, St. Louis, MO, USA) for 48 h at 37°C and 5% CO_2_. Additionally, PBMCs from A3-5 were cultured with 250 ng/mL Poly(I:C)-LMW/LyoVec™ (PolyI:C; Invivogen, San Diego, USA) for 48 h at 37°C and 5% CO_2_. Afterwards, PBMCs were carefully collected by up and down pipetting and transferred to a DNase/RNase-free tube. After centrifugation, supernatants were removed and lysis buffer for RNA extraction was added to the cell pellet. All samples in lysis buffer were stored at -80[until RNA extraction.

### RNA extraction and quantification

Total RNA was extracted from PBMCs using the RNeasy® Mini Kit (Qiagen Ltd., Crawley, UK), according to the manufacturer’s protocol. After RNA elution, an additional DNase I treatment was performed using the Heat&Run gDNA removal kit (ArcticZymes Technologies, Tromsø, Norway), following the manufacturer’s protocol. Finally, RNase inhibitors (Invitrogen, Life Technologies, Waltham, USA) were added to the RNA samples in a final concentration of 1 U/µL prior storage at -80°C until reverse transcription (RT) reaction was performed.

The purity, quantity and integrity of the extracted RNA were assessed using a BioDrop *µLITE* Spectrophotometer (BioDrop Ltd, Cambridge, UK) and Lab-Chip analysis (Agilent Technologies, Santa Clara, USA). The A260:A280 ratio ranged from 1.6 to 2.1, and RNA Integrity Numbers (RIN) ranged from 7 to 9.6.

### cDNA synthesis

Total RNA samples were used to generate cDNA as previously described (Te et al., 2021). Briefly, 110 ng of RNA were retrotranscribed in a final volume of 10 μL using the PrimeScript RT reagent Kit (Takara, Kusatsu, Japan) with a combination of oligo-d(T) and random hexamers, according to the manufacturer’s protocol. No-reverse transcription controls (no-RT) with all buffers and reagents supplied by the kits, but omitting the reverse transcriptase, were prepared to assess non-specific amplifications and presence of genomic DNA (gDNA).

Additionally, control cDNA samples from stimulated PBMCs were obtained with the aim to generate standard curves and determine primer pair efficiencies. Samples were pooled by species at the same proportion per individual animal, except for the dromedary camel. For each species, pools contained cDNA samples from PMA-ionomycin and PHA-stimulated PBMCs at 1:1 proportion, while alpaca PBMCs stimulated with PolyI:C were pooled independently. Finally, samples were serially diluted by 1:4 steps (1/20, 1/80, 1/320, 1/1280, 1/5120) prior amplification reactions.

### Primer design of immune associated and reference genes

Camelid genes and mRNA were found through bibliographic search or with described mRNAs in other species performing BLASTn (https://blast.ncbi.nlm.nih.gov/Blast.cgi). Primers were designed through comparative genomics of sequences deposited at the National Centre for Biotechnology Information (NCBI) GenBank database of llama (*Lama glama*), alpaca (*Vicugna pacos*), dromedary camel (*Camelus dromedarius*), bactrian camel (*Camelus bactrianus*), and wild bactrian camel (*Camelus ferus*). Comparison of mRNA and genomic sequences of each studied gene were performed with the alignment tool ClustalW to determine exon boundaries. In some instances, exons were already annotated in camelid genomes.

Primer pairs were designed with Primer3 (http://bioinfo.ut.ee/primer3-0.4.0/), Primer-Blast (https://www.ncbi.nlm.nih.gov/tools/primer-blast), or Primer Express 2.0 (ThermoFisher Scientific, Life Technologies, Waltham, USA), according to the following desirable criteria: (i) to span two or more exons, and some of them were placed at the exon-exon boundaries, (ii) 17-23 nucleotides in length, close to the mRNA 3’ end when possible, (iii) GC-content percentage between 45 and 55%, (iv) leading to an approximate 80-200 bp PCR product, (v) melting temperature (Tm) of each primer between 57-63°C with less than 2°C difference within primer pairs, and (vi) avoiding primer hairpin, self-primer dimer or cross-primer dimer formation.

The avoidance of primer secondary structure arrangement was assessed through the Beacon Designer™ program (http://www.premierbiosoft.com/qOligo/Oligo.jsp?PID=1), selecting for primers with ΔG greater than -3.5 kcal/mol when possible. Further, primer sequence specificity was assessed using the BLASTn alignment tool against all camelid genome sequences. Potential transcription of predicted pseudogenes was assessed by carrying out promoter region analyses through the VISTA (http://genome.lbl.gov/vista/customAlignment.shtml) and the Promoter 2.0 Prediction Server (http://www.cbs.dtu.dk/services/Promoter) softwares. Finally, the primer position within exons was checked *in silico* with their respective camelid gene sequences from the NCBI GenBank using the MapViewer tool. All the primers designed in this study are summarized in Table1. Table S1 compiles GenBank accession numbers of camelid genes and mRNA used in this study. Table S2 compiles the principal characteristics of genes and derived primers used in this study. Oligonucleotides used in this study were supplied by Roche Diagnostics (Sant Cugat del Vallès, Barcelona, Spain).

### Cytokine quantification by Fluidigm Biomark microfluidic RT-qPCR

cDNA obtained from PBMCs samples were used to validate the whole panel of primers designed for camelid species and to quantify gene expression levels by a microfluidic qPCR technique. Firstly, cDNA samples were pre-amplified using the TaqMan PreAmp Master Mix (Applied Biosystems, Life Technologies, Waltham, USA), following the manufacturer’s recommendations, doing an initial activation step of the AmpliTaq Gold DNA Polymerase for 10 min at 95°C, followed by 16 cycles of 15 seconds denaturation at 95°C plus 4 min annealing and extension at 60°C. Pre-amplified products were treated with Exonuclease I (New England Biolabs, Ipswich, USA) for 30 min at 37°C to eliminate the carryover of unincorporated primers. An inactivation step of the enzyme for 15 min at 80°C was included according to the manufacturer’s protocol. The 96.96 Dynamic Array IFCs, the 96.96 DNA Binding Dye Sample/Loading Kit (Fluidigm Corporation, South San Francisco, USA) was prepared according to the manufacturer’s instructions. Pre-amplified samples were diluted 1/20 in 1x TE Buffer, and aliquots of 2.25 µL of each sample and 0.6 µL of primer pairs at 100 µM were loaded in duplicates into their respective array inlets. Quantification of PCR reactions was performed on a Biomark HD system (Fluidigm Corporation, South San Francisco, USA). The PCR consisted in an initial activation step of 1 min at 95°C, followed by 30 cycles of 5 seconds at 96°C plus 1 min at 60°C. A dissociation step, increasing 1°C every 3 seconds from 60 to 95°C, was included for all reactions to confirm single specific PCR product amplification and define the Tm of each amplicon. Additionally, stimulated control samples were assayed in triplicates to create relative standard curves and calculate primer amplification efficiencies (see Table S3). No-RT controls and no-RNA template controls (NTC) were included in each assay to check for non-specific amplification or primer-dimer formation.

### Relative quantification and data analysis

Expression data was collected with the Fluidigm Real-Time PCR analysis software 4.1.3 (Fluidigm Corporation, South San Francisco, USA). Cycle of quantification (Cq) threshold detection value was set at 0.020, quality threshold cut-off value was established at 0.65 and amplification specificity was assessed by Tm analyses for each reaction. Amplifications fulfilling the above criteria were analysed using the DAG expression software 1.0.5.6 (Ballester et al., 2013) to apply the relative standard curve method (Livak, 1997). Cq values obtained from pooled cDNA controls were used to create standard curves for each gene, species and PBMC stimulation condition, and to extrapolate the relative quantity values. R-squared values were determined for each standard curve and the specific PCR efficiencies were calculated by applying the formula (10^(−1/slope value)-1)*100 (see Table S3). Multiple reference gene normalization was performed by using glyceraldehyde-3-phosphate dehydrogenase (*GAPDH*), hypoxanthine phosphoribosyltransferase (*HPRT1*) and ubiquitin C (*UBC*) as endogenous controls. Their suitability for normalization procedures was assessed by control-gene stability analyses with the DAG expression software 1.0.5.6 (Ballester et al., 2013). The normalized quantity values of each sample and assay were used for direct comparison in relation to non-stimulated PBMCs samples cultured during 48 h. Therefore, the up- or down-regulated expression of each gene was expressed in fold changes (Fc).

Statistical analyses could only be applied in results from alpaca PBMCs cultured with PolyI:C stimulus, due to the sample size. Fc values were logarithmically transformed to achieve normal distributions. Means of the transformed fold changes obtained for stimulated and unstimulated conditions were compared using unpaired t-test analyses in R and GraphPad Prism softwares. Differences were considered significant at *p*-values < 0.05.

## RESULTS

### Selection of immune-related genes and primer design

The selected genes encompassed several functional categories representative of pathogen innate and adaptive immune responses, and comprised type I, II and III IFNs, pattern recognition receptors (PRRs), transcription factors (TFs), IFN-stimulated genes (ISGs), pro- and anti-inflammatory cytokines, enzymes, adaptors, cellular receptors, and other genes involved in Th1 and Th2 responses. In addition, three reference genes, *GAPDH, HPRT1* and *UBC*, were selected to normalize gene expression. Table 1 summarizes genes and primers designed for subsequent expression analyses.

**Table 1.**
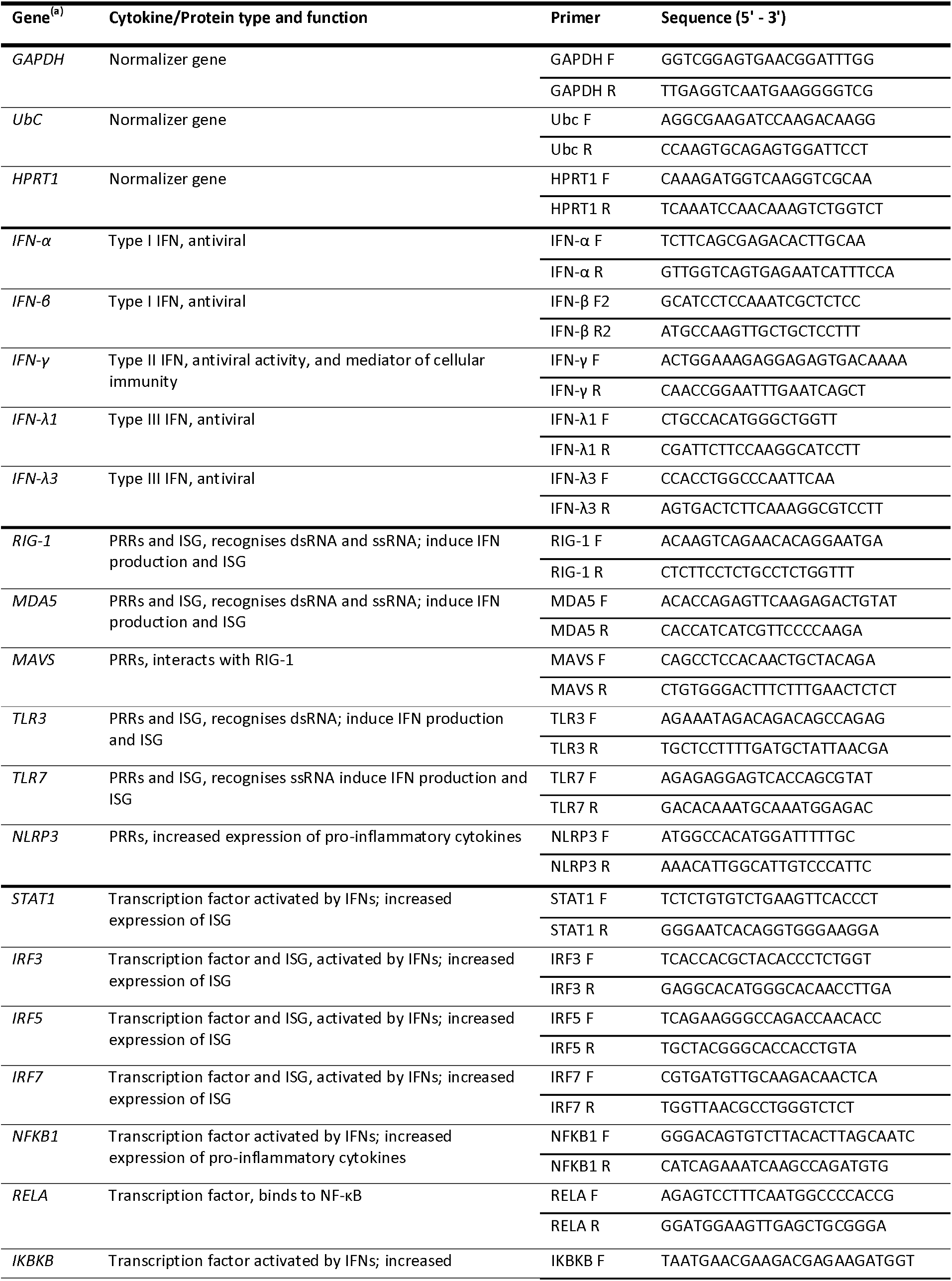
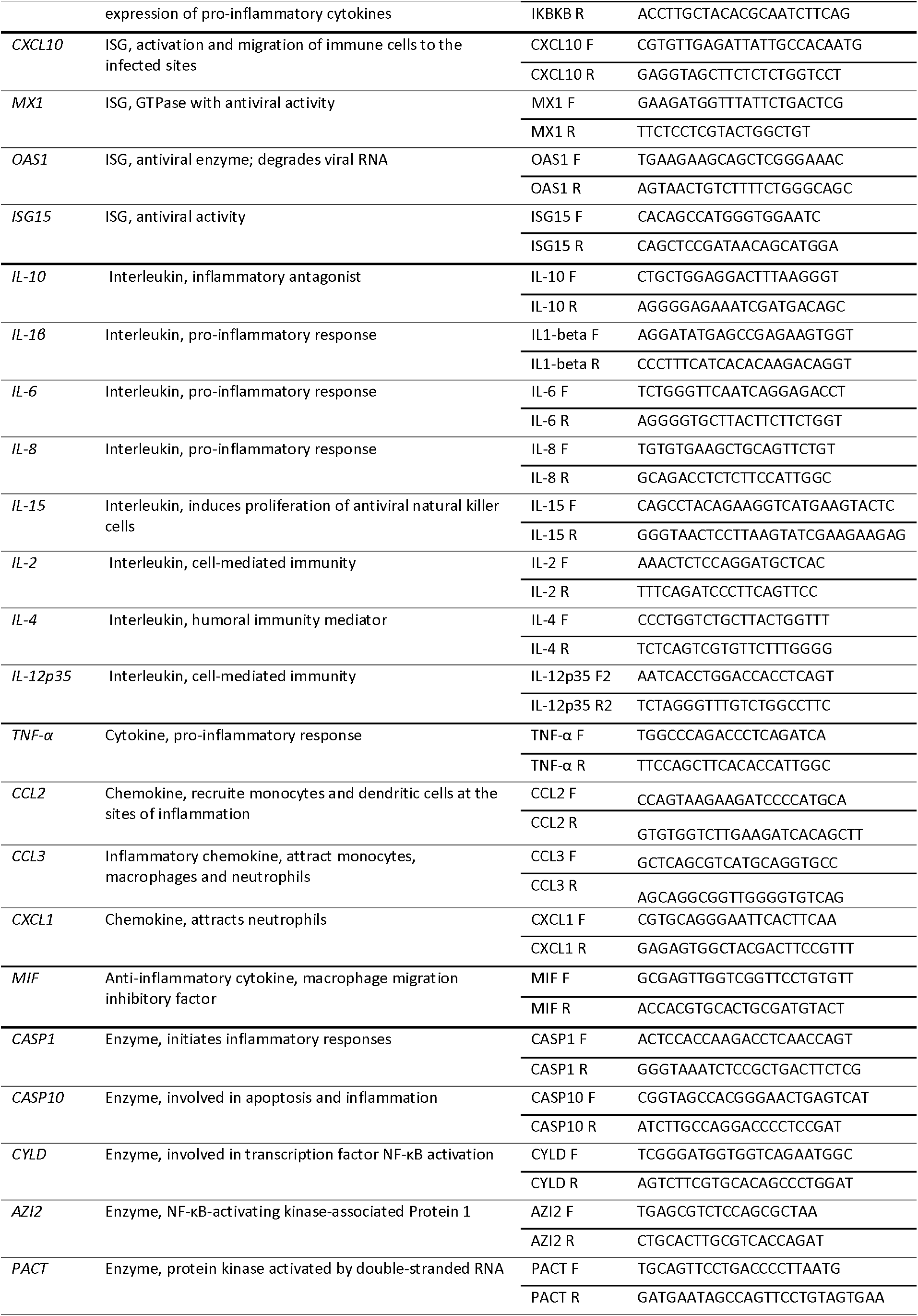
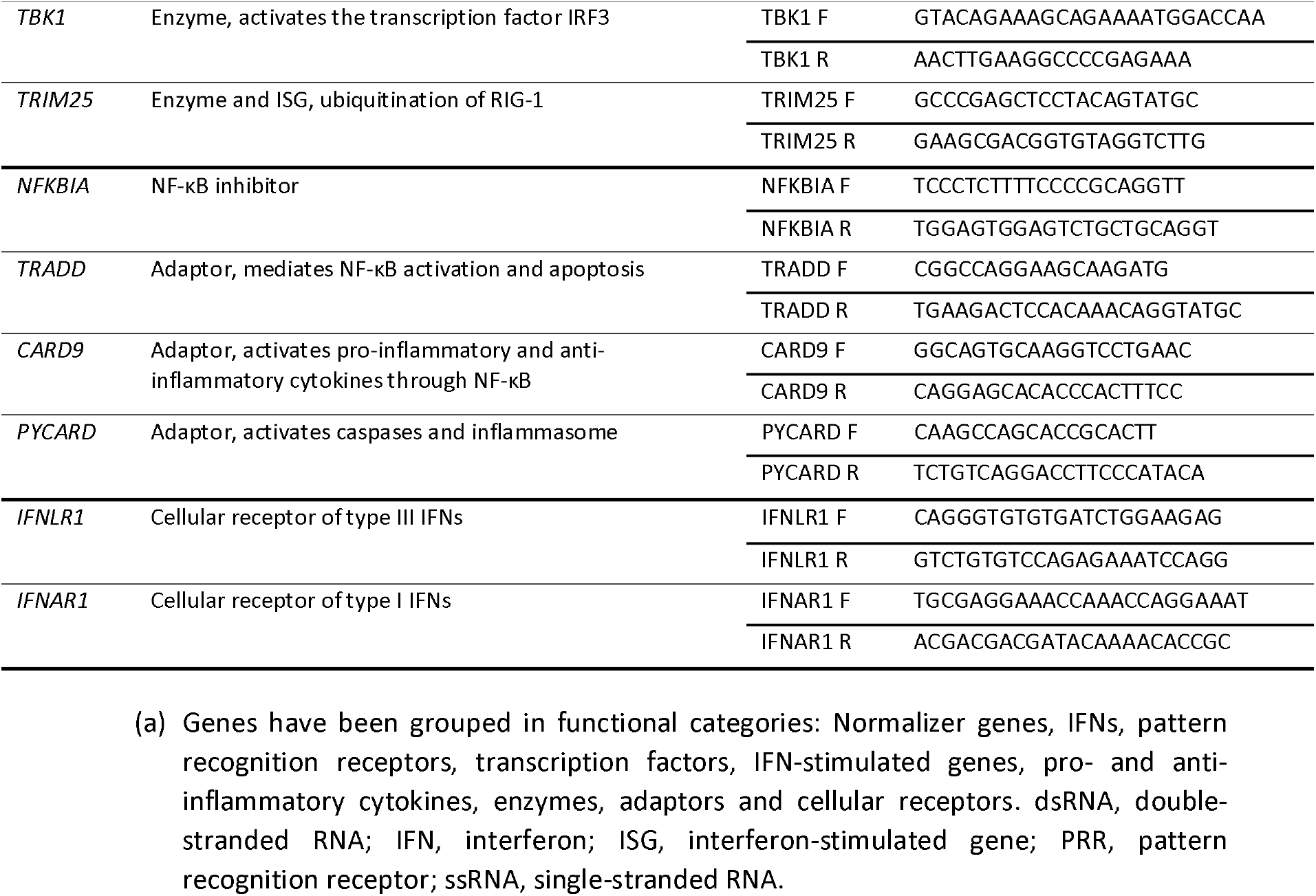
Features of the selected cytokines and immune genes used for gene expression analyses, and their validated primer pair sequences working for all camelid species.

Primer sets were designed to anneal in conserved transcribed regions of genes from five camelid species (alpaca, Bactrian camel, dromedary camel, llama, and wild Bactrian camel). Some of the genes were not annotated in the genome of llama (see Table S1) but exon/intron boundaries could be found by performing a BLASTn with the mRNA of other camelid species. The main features of the designed primer pairs are listed in Table S2.

### Primer amplification efficacy and specificity

Pools of cDNAs prepared from stimulated PBMCs were used to evaluate the whole panel of primers in samples from alpaca, dromedary, and llama. Number of dilutions used for the generation of each standard curve, slopes, coefficients of determination and amplification efficiencies are listed in Table S3. After 48 h of PBMC stimulation, all gene transcripts were sufficiently expressed to generate standard curves using 3 to 5 serial dilutions, except for *IFNAR1* in samples from dromedary and *IFN-*λ*1* in those from dromedary camel and llama (Table S3). *IFN-*λ*3* was not expressed in PBMCs from any species, regardless of the stimuli type (Table S3). All calibration curves produced linear standard curves, as evidenced by high coefficients of determination (>0.95, except for 1 sample that was 0.886). Primer pairs resulted in optimal amplification efficiencies in the different camelid species, ranging from 69 to 100%. Tm analyses confirmed a single specific amplicon for all amplified gene transcripts in all three camelid species. No amplifications occurred in no-RT and NTC samples included in the microfluidic RT-qPCR assays. Thus, most of the designed primer pairs targeting camelid cytokines and immune-related genes, as well as endogenous genes, displayed optimal specificity and efficacy suitable for immune response studies in camelid PBMCs.

### Gene expression analyses of llama PBMCs by microfluidic RT-qPCR

Transcriptomic gene expression profile of llama PBMCs stimulated for 48 h with PHA and PMA-ionomycin were compared to those of unstimulated cells. Relative expression of the different genes grouped by functional categories is shown in Figure 1a-j. PHA and PMA-ionomycin stimulation provoked similar gene expression profiles in llama PBMCs. Both stimuli induced expression of *IFN-*γ at high levels (114 and 123 Fc, respectively), but none of the other type I or III IFNs were upregulated (Figure 1a). Within the category of ISGs, *CXCL10* expression was induced by PHA and PMA-ionomycin and *MX1* only by PHA stimulated samples (Figure 1c). PHA and PMA-ionomycin also provoked the upregulation of *CCL3, TNF-*α, *IRF7* and *NFKB1* (Figure 1d, e and f). High levels of *IL-6* (Fc of 79.82 and 62.47), *IL-2* (Fc of 71.32 and 80.30) and *IL-4* (Fc of 195.78 and 145.58) expression were found of PHA and PMA-ionomycin stimulation, respectively (Figure 1e and j). Transcription of other pro- or anti-inflammatory cytokines, PRRs, adaptors, enzymes and IFN receptors were not induced in llama stimulated PBMCs (Figure 1b, e, g, h and i). Globally, the results obtained in llama cells were according to the expectancy that PHA and PMA-ionomycin provoke a marked polyclonal stimulation of PBMCs.

**Figure 1.**
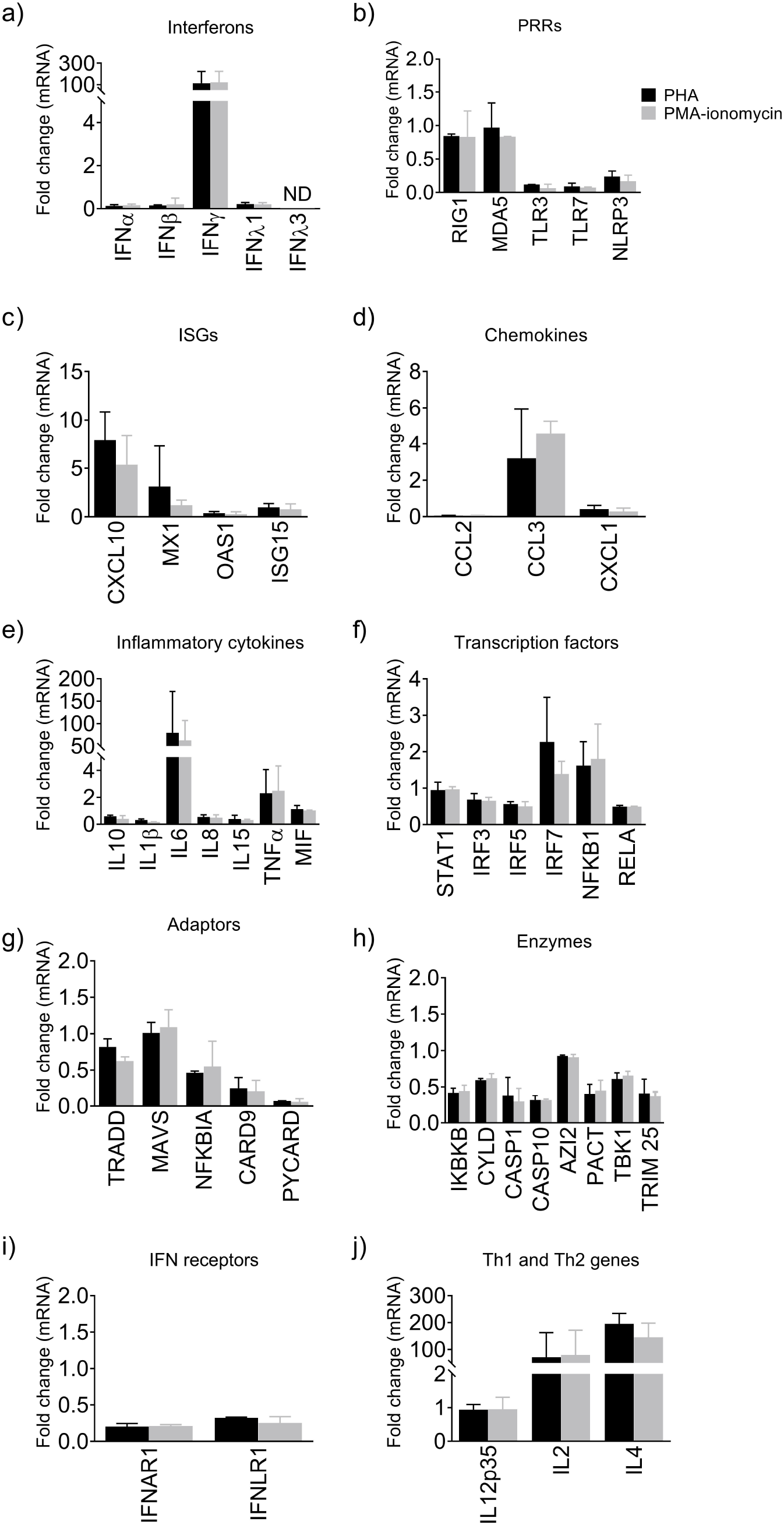
Relative expression of llama immune genes by microfluidic RT-qPCR. Gene expression profile of llama (L1-2) PBMCs stimulated for 48 h with PHA or PMA-ionomycin were compared to that from unstimulated cells. The relative standard curve method was applied for normalization purposes using multiple reference gene normalization (*GAPDH, HPRT1* and *UbC*). Immune genes were grouped by functional categories: (a) IFNs, (b) PRRs, (c) ISGs, inflammatory (d) chemokines and (e) cytokines, (f) transcription factors, downstream signalling (g) adaptors and (h) enzymes, (i) cellular receptors, and (j) cytokines involved in Th1 and Th2 response. Black and grey bars display differential expression of PHA and PMA-ionomycin stimulated PBMCs, respectively, relative to unstimulated cells. Relative expression data is displayed as mean fold-change differences ± SD. IFN, interferon; ISG, interferon-stimulated gene; PBMCs, peripheral blood mononuclear cells; PHA, phytohemagglutinin; PMA, phorbol 12-myristate 13-acetate; PRR, pattern-recognition receptor.

### Gene expression analyses of alpaca PBMCs by microfluidic RT-qPCR

We utilized the same methodology to study immune gene expression in alpaca. Gene expression profile of stimulated PBMCs (PHA, PMA-ionomycin and PolyI:C) were compared to that of unstimulated cells (Figure 2a-j). PHA and PMA-ionomycin stimulation triggered an upregulation of *RIG-1, MDA5, MX1, ISG15, CXCL1*, and *IL-8* compared to non-stimulated samples cultured for 48 h (Figure 2b, c, d, e). Moreover, expression of *IFN-*β, *IFN-*γ, *IL-10, IL-6, STAT-1, IRF7, NFKB1, CASP1, IL-2* and *IL-4* was more markedly upregulated in PHA-stimulated PBMCs (Figure 2a, e, f, h and j).

**Figure 2.**
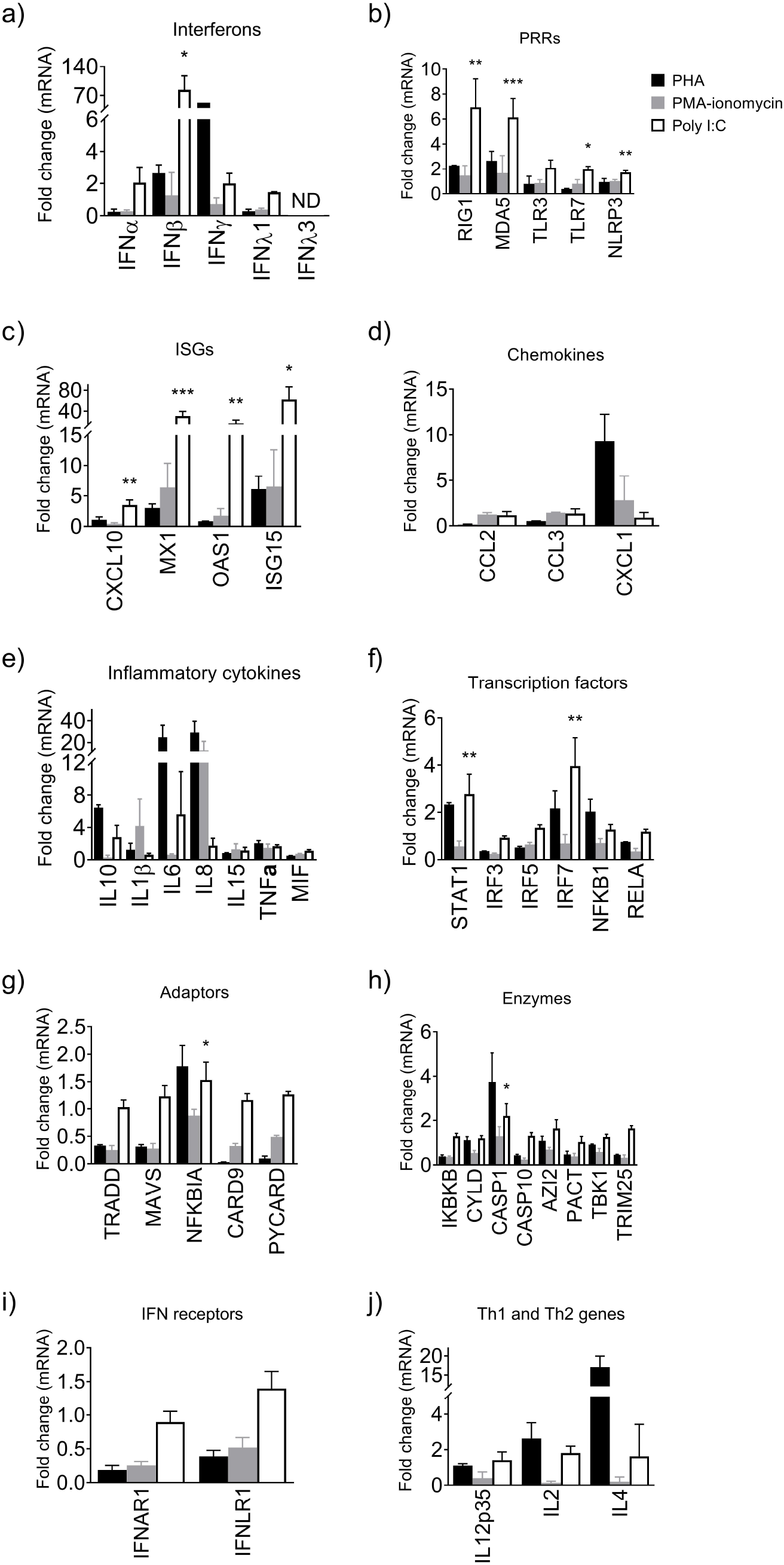
Relative expression of alpaca (A1-5) immune genes by microfluidic RT-qPCR. Immune gene expression profile of alpaca PBMCs stimulated with PHA (A1-2, black bars), PMA-ionomycin (A1-2, grey bars), or PolyI:C (A3-5, empty bars) for 48 h were compared to that from non-stimulated cells. The relative standard curve method was applied for normalization purposes using multiple reference gene normalization (*GAPDH, HPRT1* and *UbC*). Immune genes were grouped by functional categories: (a) IFNs, (b) PRRs, (c) ISGs, inflammatory (d) chemokines and (e) cytokines, (f) transcription factors, downstream signalling (g) adaptors and (h) enzymes, (i) cellular receptors, and (j) cytokines involved in Th1 and Th2 response. Relative expression data is displayed as mean fold-change differences ± SD. Statistical significance was determined by unpaired t-test. *indicates *p*-value < 0.05; **indicates *p*-value < 0.01; ***indicates *p*-value < 0.001 compared with control samples obtained prior cell culture. IFN, interferon; ISG, interferon-stimulated gene; PBMCs, peripheral blood mononuclear cells; PHA, phytohemagglutinin; PMA, phorbol 12-myristate 13-acetate; PRR, pattern-recognition receptor.

On the other hand, PMBCs from three additional alpacas were cultured with PolyI:C to ensure functionality of primers targeting mRNA of impassive genes to PHA or PMA-ionomycin stimulation. PolyI:C exposure for 48 h resulted in the upregulation of different immune genes when compared with the previous polyclonal stimulations (*IFN-*α, *IFN-*β, *RIG-1, MDA5, TLR3, TLR7, CXCL10, MX1, OAS1, ISG15, IL-10, IL-6, STAT1* and *IRF7*), some of them being expressed at high levels such as *IFN-*β (83.72 Fc), *MX1* (30.68 Fc) and *ISG15* (62.34 Fc) (Figure 2a-f). Moreover, statistical analyses determined a significant increase in *IFN-*β, *RIG-1, MDA5, TLR7, NLRP3*, all ISGs, *STAT1* and *IRF7* expression levels compared to non-stimulated alpaca PBMCs.

Therefore, these results confirmed Poly I:C as a good *in vitro* immunostimulant of antiviral responses in alpaca PBMCs.

### Gene expression analyses of dromedary PBMCs by microfluidic RT-qPCR

Expression of dromedary immune genes in stimulated PBMCs were compared to that of unstimulated cells (Figure 3a-j). Dromedary cells stimulated with PHA and PMA-ionomycin displayed similar transcriptomic profiles with characteristics of classical polyclonal stimulations. Relative to non-stimulated control samples, both PHA and PMA-ionomycin stimuli provoked the induction of *IFN-*β, *IFN-*γ, *CCL3, CXCL1, IL-6, IL-8, TNF-*α, *MIF, NFKB1, NFKBIA* and *IL-4* in higher relative expression levels than unstimulated PBMCs (Figure 3a, d, e, f, g and j). In addition, specific PHA stimulation led to increased mRNA levels of *IFN-*α, *IFN-*λ*1, RIG-1, MDA5, CXCL10, MX1, OAS1, ISG15, IL-10, IL-1*β, *IRF7* and *TRIM25*, while transcription of *IL-2* was markedly induced in PMA-ionomycin stimulated PBMCs of dromedary camels (Figure 3a, b, c, e, f, h and j). Overall, dromedary PBMCs were properly stimulated by PHA and PMA-ionomycin, enhancing immune-related gene expression in accordance with regular polyclonal stimulations.

**Figure 3.**
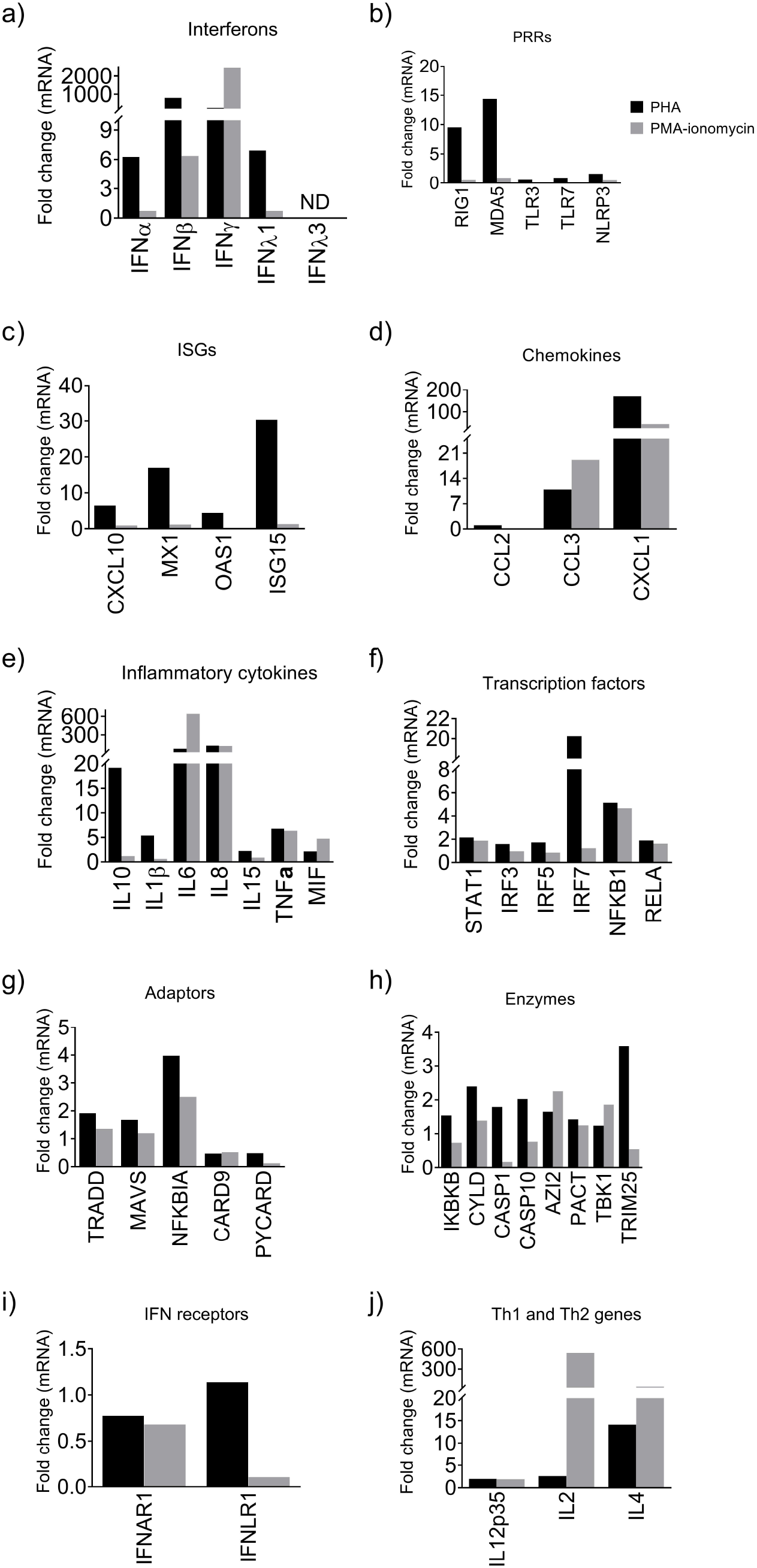
Relative expression of dromedary camel (D1) immune genes by microfluidic RT-qPCR. Immune gene expression profile of dromedary PBMCs stimulated with PHA (black bars) or PMA-ionomycin (grey bars) for 48 h were compared to that from non-stimulated cells. The relative standard curve method was applied for normalization purposes using multiple reference gene normalization (*GAPDH, HPRT1* and *UbC*). Immune genes were grouped by functional categories: (a) IFNs, (b) PRRs, (c) ISGs, inflammatory (d) chemokines and (e) cytokines, (f) transcription factors, downstream signalling (g) adaptors and (h) enzymes, (i) cellular receptors, and (j) cytokines involved in Th1 and Th2 response. Relative expression data is displayed as mean fold-change differences ± SD. Statistical significance was determined by unpaired t-test. *indicates *p*-value < 0.05; **indicates *p*-value < 0.01; ***indicates *p*-value < 0.001 compared with control samples obtained prior cell culture. IFN, interferon; ISG, interferon-stimulated gene; PBMCs, peripheral blood mononuclear cells; PHA, phytohemagglutinin; PMA, phorbol 12-myristate 13-acetate; PRR, pattern-recognition receptor.

## DISCUSSION

Due to the lack of reagent availability (i.e., antibodies) in several animal species, analysis of gene expression has become a common method to determine the immune transcriptomic profile after infection and/or vaccination. In this work, we implemented an RT-qPCR method to monitor camelid innate and adaptive immune responses, which allow the quantification of forty-seven cytokines and immune-related genes in a single run. Importantly, the designed primer sets can be used for all camelid species using the same conditions (primer hybridization temperature) and methodology.

Several infectious diseases affect camelid species and threaten livestock productivity (Camel pox virus, trypanosomiasis, and gastro-intestinal helminthiases, among others). Also, camelids are a source for several zoonotic viral and bacterial pathogens (MERS-CoV, Crimean-Congo haemorrhagic fever virus, *Rickettsia* spp, and others) which threaten public health (Khalafalla, 2017; Wernery et al., 2014). However, immune responses elicited upon infections of camelids remain largely unknown. Although a previous work provided tools to quantify inflammatory cytokines mRNA from llama (Odbileg, Konnai, et al., 2004), the primers were designed before available annotation of camelid genomes (Wu et al., 2014). Therefore, we designed new primers sets to perform more accurate assays.

First, we focused on the requirements to ensure a specific amplification of the products (Taylor et al., 2010) and established criteria for the subsequent design of primers. Although not all primer designs fulfilled all parameters of the criteria (i.e., genes with a single exon as IFNs α, β, and λ1), specific amplification was determined together with the absence of amplification in no-RT and NTC controls, which ensured that gDNA amplification did not occur. In addition, three reference genes, (*GAPDH, HPRT1* and *UBC*) were selected to normalize gene expression (Cicinnati et al., 2008; Mehta et al., 2010; Souza et al., 2013). These genes have been reported to be stable for data normalization in T lymphocytes (Albershardt et al., 2012; Bas et al., 2004; Dheda et al., 2004) and respiratory samples (Resa et al., 2014; Te et al., 2021). Choosing appropriate reference genes is crucial to achieve optimal data normalization, which is mandatory to discard sample-to-sample variations. Here, the selected normalizer genes were stable in PBMC samples from all the studied species, regardless the type of stimulation.

The designed primers would allow studying camelid immune responses in most laboratories worldwide, including those in developing countries, which have been infrastructurally and technically upgraded to perform PCR-based assays since the recent influenza virus outbreaks and the COVID-19 pandemics (Breiman et al., 2007). Nonetheless, to study expression analyses of a broad panel of genes in multiple samples by the gold standard RT-qPCR can result tedious and relatively expensive. Thus, we integrated the whole set of designed primers in a unique Fluidigm Biomark microfluidic qPCR assay. Gene expression analyses showed that PHA and PMA-ionomycin stimulation produced strong induction of some cytokines and immune gene transcripts in camelid PBMCs, which displayed profiles commonly found in polyclonally activated lymphocytes of bovine (Norian et al., 2015) and human (Rostaing et al., 1999). PBMCs from dromedary camel and llama underwent a robust increase in *IFN-*γ, *IL-4* and *IL-6* expression after PHA and PMA-ionomycin exposure, and the same phenomenon occurred in alpaca PBMCs stimulated with PHA but not with PMA-ionomycin. Transcription of the pro-inflammatory *IL-8* and the anti-inflammatory *IL-10* was also upregulated in polyclonally-stimulated alpaca and dromedary cells, but not in those from llama. Upregulation of *IL-8* in camelid cells was expected since it is tightly regulated upon activation of the transcription factor NF-κB (Liu et al., 2017). Similar to previous results in humans (Baran et al., 2001), *IL-10* expression was only upregulated by PHA stimulation but PMA-ionomycin was ineffective. Furthermore, it is known that type I and III IFNs activate a different signalling pathway than *IFN-*γ does, which can lead to the expression of distinct ISGs (Sen et al., 2018; W. Wang et al., 2017). *CXCL10* and *CCL3* are typically classified as ISGs induced by *IFN-*γ (Rabin, 2003; Sen et al., 2018). Upregulation of these chemokines was observed in all dromedary and llama samples expressing high levels of *IFN-*γ, apart from *CXCL10* in dromedary cells stimulated with PMA-ionomycin. As reported here, the stimulation of camelid cells using PHA and PMA-ionomycin did not activate the transcription of PRRs, adaptors, enzymes and IFN receptors compared to unstimulated controls. As observed in other mammalian species (Ciliberti et al., 2017; Gao et al., 2010; Lin et al., 2021; Norian et al., 2015; Wagner & Freer, 2009), PHA and PMA-ionomycin are also potent antigen surrogate activators of Th1 and Th2 cytokine expression in alpaca, dromedary, and llama PBMCs. Our results support that PHA and PMA-ionomycin can be used to monitor camelid T-cell activation, proliferation, and effective cytokine production.

Curiously, a broader gene expression profile was observed in dromedary PBMCs 48 h after PHA stimulation. Besides the activation of type II IFN signalling pathway, PHA also upregulated the expression of type I and III IFNs, as well as downstream genes regulated upon the IFN signalling cascade activation. Consequently, strong induction of PRRs and ISGs occurred in dromedary samples but not in those from alpaca or llama. Expression of *IRF7* and *IL-10* were also exclusively upregulated in PHA-stimulated dromedary PBMCs, in agreement with previous reports showing that both type I and III IFNs increase the transcription of *IRF7* factor *in vitro* (Zhou et al., 2007), and that IL-10 production is enhanced as a type III IFN-stimulated gene (Stanifer et al., 2019).

Further analyses involving samples from more animals would help to rule out individual animal variations to confirm whether dromedary camels have a broader immune response after PHA stimulation than other camelid species.

*In vitro* stimulation of PBMCs with PolyI:C is known to elicit the expression of cytokines, mimicking certain aspects of viral infection (Huang et al., 2006). After 48 h stimulation, alpaca PBMCs sensed PolyI:C and increased the expression of type I IFNs, all studied PRRs and ISGs, along with the transcription factors *STAT1* and *IRF7*. Contrary to human PBMCs (Murata et al., 2014), alpaca lymphocytes exposed to PolyI:C induced type I but not type III IFNs. Previous investigations indicated that *IFN-*α was not produced by ovine and bovine PBMCs treated with PolyI:C (Booth et al., 2010), hence future studies could shed light on differential antiviral immune responses among livestock species. Globally, PolyI:C stimulation of alpaca PBMCs yielded similar results than porcine and human PBMCs did (Docampo et al., 2017; Huang et al., 2006; J. Wang et al., 2016). Thus, Poly I:C is a good immunostimulant of antiviral responses in alpaca PBMCs. Further research is needed to determine if other camelid species are equally responsive to Poly I:C.

In the current work, regardless of the stimuli used to boost cytokine expression, none of the PBMC stimulation assays in any camelid species showed detectable levels of *IFN-* λ*3* expression. However, we previously demonstrated an optimal amplification of *IFN-* λ*3* mRNA in the nasal epithelium of MERS-CoV infected alpacas (Te et al., 2021, 2022) using the pair of primers reported here. Primers targeting *IFN-*λ*3* were also designed to anneal mRNA of all camelid species, therefore, a correct amplification is also expected in samples from other camelids, although further studies are needed to confirm this hypothesis.

In summary, we implemented a RT-qPCR method for the simultaneous quantification of cytokines and immune-related genes involved in major immune response signalling pathways of different camelid species. The novel assay was set up after the design of primers targeting immune genes and performing data normalization with three reference genes. The assays were validated using PBMCs from alpaca, dromedary camel and llama PBMCs after stimulation with PHA, PMA-ionomycin or PolyI:C. Microfluidic RT-qPCR results indicated that PBMCs from all camelid species stimulated with PHA and PMA-ionomycin induce genes characteristic of Th1 and Th2 responses, besides PHA activation of type I and III IFN signalling pathways in dromedary lymphocytes. PolyI:C stimulation produced a marked antiviral response in alpaca PBMCs.

## ACKNOWLEDGEMENTS

The authors thank Joana Ribes from the Centre for Research in Agricultural Genomics (CRAG, Cerdanyola del Vallès, Barcelona) for her technical guidance in the use of Fluidigm BioMark technology. This study was performed as part of the Zoonotic Anticipation and Preparedness Initiative (ZAPI project) [Innovative Medicines initiative (IMI) grant 115760] and the Veterinary Biocontained research facility Network (VetBioNet) project (EU Grant Agreement INFRA-2016-1 Nº731014), with assistance and financial support from IMI and the European Commission and contributions from EFPIA partners. J.R. was partially supported by the VetBioNet project. IRTA is supported by CERCA Programme / Generalitat de Catalunya.

## AUTHOR CONTRIBUTIONS

J.R., J.S., J.V.-A. and A.B. conceived and designed the study. J.R., N.T., M.B., A.B., J.S., J.V.-A and A.B. performed the experiments and analyzed the data. All the authors discussed the results. The manuscript was written by J.R. and all the authors revised the manuscript.

---

**Table S1**. NCBI accession numbers of camelid gene sequences used for comparative analyses and primer design.

**Table S2**. Features of the primer pairs designed for the quantification of camelid immune and reference genes by RT-qPCR.

**Table S3**. Performance of the primer pairs designed for the quantification of mRNA expression of camelid immune and reference genes.

**Table S4**. Expression level of immune response genes from alpaca, dromedary camel and llama PBMCs stimulated with PHA, PMA-ionomycin or PolyI:C.

