## Supplementary material for "High-throughput quantification of camelid cytokine mRNA expression in PBMCs by microfluidic qPCR technology": Table S1

**Table S1**. NCBI accession numbers of camelid gene sequences used for comparative analyses and primer design.

| Gene | Species | GenBank accession number |
|---|---|---|
| *GAPDH* | Camelus dromedarius | XM_010990867.2 |
| | Camelus bactrianus | XM_010957730.1 |
| | Vicugna pacos | XM_006210852.2 |
| | Camelus ferus | XM_006181646.3 |
| *HPRT1* | Camelus dromedarius | XM_031446174.1 |
| | Camelus bactrianus | XM_010968460.1 |
| | Vicugna pacos | XM_031671409.1 |
| | Camelus ferus | XM_032474943.1 |
| *UbC* | Camelus dromedarius | XM_031442494.1 |
| | Camelus bactrianus | XM_010969735.1 |
| | Vicugna pacos | XM_031670203.1 |
| | Camelus ferus | XM_032471921.1 |
| *IFN-α* | Camelus dromedarius | XM_010999340.2 |
| | Camelus bactrianus | XM_010946010.2 |
| | Vicugna pacos | XM_015242649.2 |
| | Camelus ferus | XM_032477607.1 |
| *IFN-β* | Camelus dromedarius | XM_010988144.1 |
| | Camelus bactrianus | XM_010958977.2 |
| | Vicugna pacos | XM_006208258.1 |
| | Camelus ferus | XM_006180372.1 |
| *IFN-γ* | Lama glama | AB107652.1 |
| | Camelus dromedarius | XM_031462226.1 |
| | Camelus bactrianus | XM_010970501.1 |
| | Vicugna pacos | XM_006205835.2 |
| | Camelus ferus | XM_006189690.2 |
| *IFN-λ1* | Camelus dromedarius | XM_010978654.2 |
| | Camelus bactrianus | XM_010958621.1 |
| | Vicugna pacos | XM_006206664.2 |
| | Camelus ferus | XM_006174985.2 |
| *IFN-λ3* | Camelus dromedarius | XM_010985161.1 |
| | Camelus bactrianus | XM_010947318.1 |
| | Vicugna pacos | XM_006219677.1 |
| | Camelus ferus | XM_006195351.1 |
| *RIG-1 (DDX58)* | Camelus dromedarius | XM_010975810.2 |
| | Camelus bactrianus | XM_010967358.2 |
| | Vicugna pacos | XM_031676595.1 |
| | Camelus ferus | XM_006192497.3 |
| *MDA5 (IFIH1)* | Camelus dromedarius | XM_010985569.2 |
| | Camelus bactrianus | XM_010971381.1 |
| | Vicugna pacos | XM_006196223.3 |
| | Camelus ferus | XM_006190090.2 |

| | | |
|---|---|---|
| *MAVS (VISA)* | Camelus dromedarius | XM_010988239.2 |
| | Camelus bactrianus | XM_010972744.1 |
| | Vicugna pacos | XM_006207415.3 |
| | Camelus ferus | XM_006187131.3 |
| *TLR3* | Camelus dromedarius | XM_010995734.2 |
| | Camelus bactrianus | XM_010953279.2 |
| | Vicugna pacos | XM_015249164.2 |
| | Camelus ferus | XM_014553913.2 |
| *TLR7* | Camelus dromedarius | XM_010993639.2 |
| | Camelus bactrianus | XM_010966214.1 |
| | Vicugna pacos | XM_006212620.2 |
| | Camelus ferus | XM_006193069.2 |
| *NLRP3* | Camelus dromedarius | XM_010997883.2 |
| | Camelus bactrianus | XM_010950280.2 |
| | Vicugna pacos | XM_031673306.1 |
| | Camelus ferus | XM_006177988.3 |
| *STAT1* | Camelus dromedarius | XM_010979711.2 |
| | Camelus bactrianus | XM_010948718.2 |
| | Vicugna pacos | XM_031678037.1 |
| | Camelus ferus | XM_006186813.3 |
| *IRF3* | Camelus dromedarius | XM_010993178.2 |
| | Camelus bactrianus | XM_045510710.1 |
| | Vicugna pacos | XM_006208451.3 |
| | Camelus ferus | XM_006173792.3 |
| *IRF5* | Camelus dromedarius | XM_010975643.2 |
| | Camelus bactrianus | XM_010947554.2 |
| | Vicugna pacos | XM_006202276.3 |
| | Camelus ferus | XM_032483311.1 |
| *IRF7* | Camelus dromedarius | XM_031448349.1 |
| | Camelus bactrianus | XM_010956145.2 |
| | Vicugna pacos | XM_015251986.1 |
| | Camelus ferus | XM_032490347.1 |
| *NFKB1* | Camelus dromedarius | XM_010980636.2 |
| | Camelus bactrianus | XM_010953589.2 |
| | Vicugna pacos | XM_031690344.1 |
| | Camelus ferus | XM_006188020.3 |
| *RELA* | Camelus dromedarius | XM_031448012.1 |
| | Camelus bactrianus | XM_010957140.2 |
| | Vicugna pacos | XM_031690897.1 |
| | Camelus ferus | XM_032489838.1 |
| *IKBKB* | Camelus dromedarius | XM_031440029.1 |
| | Camelus bactrianus | XM_010957873.2 |
| | Vicugna pacos | XM_031691563.1 |
| | Camelus ferus | XM_006188385.3 |
| *CXCL10* | Camelus dromedarius | XM_010983050.2 |
| | Camelus bactrianus | XM_010969313.2 |

| | | |
|---|---|---|
| | Vicugna pacos | XM_006198241.3 |
| | Camelus ferus | XM_006176316.3 |
| *MX1* | Camelus dromedarius | XM_031459860.1 |
| | Camelus bactrianus | XM_010958347.2 |
| | Vicugna pacos | XM_006204960 |
| | Camelus ferus | XM_032461929.1 |
| *OAS1* | Camelus dromedarius | XM_031443284.1 |
| | Camelus bactrianus | XM_010969608.2 |
| | Vicugna pacos | XM_031670190.1 |
| | Camelus ferus | XM_032472237.1 |
| *ISG15* | Camelus dromedarius | XM_010999398.2 |
| | Camelus bactrianus | XM_010957998.2 |
| | Vicugna pacos | XM_015237784.2 |
| | Camelus ferus | XM_014551219.2 |
| *IL-10* | Lama glama | AB107649.1 |
| | Camelus dromedarius | JQ917916.1 |
| | Camelus bactrianus | NM_001303520.1 |
| | Vicugna pacos | XM_006215461.3 |
| | Camelus ferus | XM_006182265.3 |
| *IL-1β* | Lama glama | AB107644.1 |
| | Camelus dromedarius | XM_010984994.2 |
| | Camelus bactrianus | XM_010958977.2 |
| | Vicugna pacos | XM_006203828.3 |
| | Camelus ferus | XM_006183589.3 |
| *IL-6* | Lama glama | AB107647.1 |
| | Camelus dromedarius | XM_010987177.2 |
| | Camelus bactrianus | AB107656.1 |
| | Vicugna pacos | XM_006201793.2 |
| | Camelus ferus | XM_006179204.2 |
| *IL-8 (CXCL8)* | Camelus dromedarius | KF843702.1 |
| | Camelus bactrianus | XM_010969343.2 |
| | Vicugna pacos | XM_006212530.3 |
| | Camelus ferus | XM_006188697.3 |
| *IL-15* | Camelus dromedarius | XM_010978726.2 |
| | Camelus bactrianus | XM_010954603.2 |
| | Vicugna pacos | XM_015249496.2 |
| | Camelus ferus | XM_006193545.3 |
| *IL-2* | Lama glama | AB107651.1 |
| | Camelus dromedarius | NM_001303548.1 |
| | Camelus bactrianus | AB246671.1 |
| | Vicugna pacos | KM205215.1 |
| | Camelus ferus | XM_006180708.3 |
| *IL-4* | Lama glama | AB107648.1 |
| | Camelus dromedarius | HM051106.1 |
| | Camelus bactrianus | AB246673.1 |
| | Vicugna pacos | XM_006212826.3 |

| | | |
|---|---|---|
| | Camelus ferus | XM_006179596.2 |
| *IL-12p35* | Lama glama | AB107653.1 |
| | Camelus dromedarius | XM_010986258.2 |
| | Camelus bactrianus | AB246672.1 |
| | Vicugna pacos | XM_031679452.1 |
| | Camelus ferus | XM_006190436.3 |
| *TNF-α* | Lama glama | AB107646.1 |
| | Camelus dromedarius | NM_001319880.1 |
| | Camelus bactrianus | NM_001319779.1 |
| | Vicugna pacos | XM_006215316.2 |
| | Camelus ferus | XM_006178751.3 |
| *MCP-1 (CCL2)* | Camelus dromedarius | XM_010979035.2 |
| | Camelus bactrianus | XM_010970431.2 |
| | Vicugna pacos | XM_006212021.3 |
| | Camelus ferus | XM_006185837.3 |
| *MIP-1α (CCL3)* | Camelus dromedarius | XM_010990500.2 |
| | Camelus bactrianus | XM_010949170.2 |
| | Vicugna pacos | XM_006213334.3 |
| | Camelus ferus | XM_006174846.3 |
| *CXCL1* | Camelus dromedarius | XM_031462577.1 |
| | Camelus bactrianus | XM_010969410.2 |
| | Vicugna pacos | XM_031684028.1 |
| | Camelus ferus | XM_032458086.1 |
| *MIF* | Camelus dromedarius | XM_031442393.1 |
| | Camelus bactrianus | XM_010955454.2 |
| | Vicugna pacos | NM_001287197.1 |
| | Camelus ferus | XM_014552697.2 |
| *CASP1* | Camelus dromedarius | XM_010993435.2 |
| | Camelus bactrianus | XM_010953176.2 |
| | Vicugna pacos | XM_015249739.2 |
| | Camelus ferus | XM_014560469.2 |
| *CASP10* | Camelus dromedarius | XM_010991974.2 |
| | Camelus bactrianus | XM_010971860.2 |
| | Vicugna pacos | XM_006205263.3 |
| | Camelus ferus | XM_006189972.3 |
| *CYLD* | Camelus dromedarius | XM_031458461.1 |
| | Camelus bactrianus | XM_010961754.2 |
| | Vicugna pacos | XM_015242273.2 |
| | Camelus ferus | XM_006183539.3 |
| *AZI2* | Camelus dromedarius | XM_010977820.2 |
| | Camelus bactrianus | XM_010959645.2 |
| | Vicugna pacos | XM_006200749.3 |
| | Camelus ferus | XM_032459400.1 |
| *PACT (PRKRA)* | Camelus dromedarius | XM_010991356.2 |
| | Camelus bactrianus | XM_010959971.2 |
| | Vicugna pacos | XM_006210217.3 |

| | | |
|---|---|---|
| | Camelus ferus | XM_032479782.1 |
| *TBK1* | Camelus dromedarius | XM_031462774.1 |
| | Camelus bactrianus | XM_010970514.2 |
| | Vicugna pacos | XM_031683111.1 |
| | Camelus ferus | XM_032493328.1 |
| *TRIM25* | Camelus dromedarius | XM_010990378.2 |
| | Camelus bactrianus | XM_010951086.2 |
| | Vicugna pacos | XM_031685141.1 |
| | Camelus ferus | XM_014556519.2 |
| *NFKBIA* | Camelus dromedarius | XM_010983796.2 |
| | Camelus bactrianus | XM_010967860.2 |
| | Vicugna pacos | XM_031678782.1 |
| | Camelus ferus | XM_032481888.1 |
| *TRADD* | Camelus dromedarius | XM_031458299.1 |
| | Camelus bactrianus | XM_010962447.2 |
| | Vicugna pacos | XM_006203676.3 |
| | Camelus ferus | XM_032486508.1 |
| *CARD9* | Camelus dromedarius | XM_031450838.1 |
| | Camelus bactrianus | XM_010955930.2 |
| | Vicugna pacos | XM_006218359.3 |
| | Camelus ferus | XM_032478794.1 |
| *PYCARD* | Camelus dromedarius | XM_010980820.2 |
| | Camelus bactrianus | XM_010972413.2 |
| | Vicugna pacos | XM_015236916.2 |
| | Camelus ferus | XM_006181480.3 |
| *IFNLR1* | Camelus dromedarius | XM_010989801.2 |
| | Camelus bactrianus | XM_045518428.1 |
| | Vicugna pacos | XM_031683805.1 |
| | Camelus ferus | XM_032495478.1 |
| *IFNAR1* | Camelus dromedarius | XM_010981032.2 |
| | Camelus bactrianus | XM_010956566.2 |
| | Vicugna pacos | XM_006216038.3 |
| | Camelus ferus | XM_014561608.2 |
