## Supplementary material for "High-throughput quantification of camelid cytokine mRNA expression in PBMCs by microfluidic qPCR technology": Table S2

**Table S2**. Features of the primer pairs designed for the quantification of camelid immune and reference genes by RT-qPCR.

| Gene name | Primer Name | Primers (5' - 3') | Exon location[(a)] | Lenght (bp) | Tm (°C) | GC% | GC Clamp | Cross Dimer (ΔG) | Self Dimer (ΔG) | Hairpin (ΔG) | Product size (bp) |
|---|---|---|---|---|---|---|---|---|---|---|---|
| *GAPDH* | GAPDH F | GGTCGGAGTGAACGGATTTGG | 2 (71%) /3 (29%) | 21 | 58.53 | 57.14 | 2 | -0.9 | 0.0 | 0.0 | 108 |
| | GAPDH R | TTGAGGTCAATGAAGGGGTCG | 3 | 21 | 57.19 | 52.38 | 2 | -0.9 | -2.0 | -0.7 | |
| *UbC* | UbC F | AGGCGAAGATCCAAGACAAGG | 2 | 21 | 57.27 | 52.38 | 2 | -3.8 | -2.0 | 0.0 | 129 |
| | UbC R | CCAAGTGCAGAGTGGATTCCT | 2 | 21 | 57.18 | 52.38 | 2 | -3.8 | -3.4 | -1.5 | |
| *HPRT1* | HPRT1 F | CAAAGATGGTCAAGGTCGCAA | 6 | 21 | 56.37 | 47.62 | 3 | -1.5 | 0.0 | 0.0 | 82 |
| | HPRT1 R | TCAAATCCAACAAAGTCTGGTCT | 7 (43%) /8 (57%) | 23 | 55.85 | 39.13 | 3 | -1.5 | 0.0 | -1.5 | |
| *IFN-α* | IFN-α F | TCTTCAGCGAGACACTTGCAA | 1 | 21 | 57.43 | 47.62 | 2 | -2.5 | -5.7 | -0.7 | 87 |
| | IFN-α R | GTTGGTCAGTGAGAATCATTTCCA | 1 | 24 | 57.1 | 41.67 | 2 | -2.5 | -1.5 | -1.5 | |
| *IFN-β* | IFN-β F2 | GCATCCTCCAAATCGCTCTCC | 1 | 21 | 58.39 | 57.14 | 2 | -2.9 | 0 | 0.0 | 99 |
| | IFN-β R2 | ATGCCAAGTTGCTGCTCCTTT | 1 | 21 | 58.27 | 47.62 | 2 | -2.9 | -0.7 | -0.5 | |
| *IFN-γ* | IFN-γ F | ACTGGAAAGAGGAGAGTGACAAAA | 3 | 24 | 57.67 | 41.67 | 1 | -1.8 | -0.8 | -0.8 | 199 |
| | IFN-γ R | CAACCGGAATTTGAATCAGCT | 3 (80%) /4 (20%) | 21 | 54.39 | 42.86 | 1 | -1.8 | -4.3 | -0.7 | |
| *IFN-λ1* | IFN-λ1 F | CTGCCACATGGGCTGGTT | 1 | 18 | 56.6 | 61.11 | 2 | -3.9 | -2.4 | -2.4 | 82 |
| | IFN-λ1 R | CGATTCTTCCAAGGCATCCTT | 1 | 21 | 55.57 | 47.62 | 2 | -3.9 | -2.4 | -2.4 | |
| *IFN-λ3* | IFN-λ3 F | CCACCTGGCCCAATTCAA | 1 | 18 | 53.77 | 55.56 | 1 | -2.4 | -4.4 | 0.0 | 81 |
| | IFN-λ3 R | AGTGACTCTTCAAAGGCGTCCTT | 1 (52.2%) / 2 (47.8%) | 23 | 59.71 | 47.83 | 1 | -2.4 | -2.4 | -2.4 | |
| *RIG-1* | RIG-1 F | ACAAGTCAGAACACAGGAATGA | 15 (73%) /16 (27%) | 22 | 55.01 | 40.91 | 1 | -3.7 | -0.9 | -0.9 | 199 |
| | RIG-1 R | CTCTTCCTCTGCCTCTGGTTT | 16 (43%) /17 (57%) | 21 | 56.55 | 52.38 | 1 | -3.7 | 0.0 | 0.0 | |
| *MDA5* | MDA5 F | ACACCAGAGTTCAAGAGACTGTAT | 14 (60%) /15 (40%) | 24 | 57.00 | 41.67 | 1 | 0.0 | -1.1 | -1.1 | 129 |
| | MDA5 R | CACCATCATCGTTCCCCAAGA | 15 (5%) /16 (95%) | 21 | 57.26 | 52.38 | 1 | 0.0 | 0.0 | 0.0 | |
| *MAVS* | MAVS F | CAGCCTCCACAACTGCTACAGA | 4 | 22 | 59.68 | 54.55 | 1 | -4.7 | -1.8 | -1.8 | 106 |
| | MAVS R | CTGTGGGACTTTCTTTGAACTCTCT | 4 (16%) / 5 (84%) | 25 | 58.73 | 44 | 1 | -4.7 | -0.6 | -0.6 | |
| *TLR3* | TLR3 F | AGAAATAGACAGACAGCCAGAG | 5 | 22 | 54.72 | 45.45 | 1 | -2.2 | 0.0 | 0.0 | 197 |
| | TLR3 R | TGCTCCTTTTGATGCTATTAACGA | 5 | 24 | 56.69 | 37.5 | 1 | -2.2 | -0.8 | 0.0 | |

| | | | | | | | | | | | |
|---|---|---|---|---|---|---|---|---|---|---|---|
| *TLR7* | TLR7 F | AGAGAGGAGTCACCAGCGTAT | 3 | 21 | 57.23 | 52.38 | 2 | -1.5 | 0.0 | 0.0 | 104 |
| | TLR7 R | GACACAAATGCAAATGGAGAC | 3 | 21 | 53.34 | 42.86 | 2 | -1.5 | -3.4 | 0.0 | |
| *NLRP3* | NLRP3 F | ATGGCCACATGGATTTTTGC | 1 | 20 | 54.55 | 45 | 2 | -3.9 | -7.2 | -0.5 | 91 |
| | NLRP3 R | AAACATTGGCATTGTCCCATTC | 1 (31.8%) / 2 (68.2%) | 22 | 55.51 | 40.91 | 2 | -3.9 | -1.5 | -1.5 | |
| *STAT1* | STAT1 F | TCTCTGTGTCTGAAGTTCACCCT | 25 (65%) /26 (35%) | 23 | 58.56 | 47.83 | 3 | -4.3 | -2.0 | -1.3 | 191 |
| | STAT1 R | GGGAATCACAGGTGGGAAGGA | 27 | 21 | 58.59 | 57.14 | 3 | -4.3 | -1.3 | 0.0 | |
| *IRF3* | IRF3 F | TCACCACGCTACACCCTCTGGT | 7 | 22 | 62.69 | 59.09 | 2 | -2.9 | -2.9 | -2.9 | 102 |
| | IRF3 R | GAGGCACATGGGCACAACCTTGA | 7 (17.4%) / 8 (86.6%) | 23 | 63.25 | 56.52 | 2 | -2.9 | -2.3 | -1.3 | |
| *IRF5* | IRF5 F | TCAGAAGGGCCAGACCAACACC | 7 | 22 | 61.84 | 59.09 | 2 | -2.4 | -4.4 | 0.0 | 121 |
| | IRF5 R | TGCTACGGGCACCACCTGTA | 7(20%) / 8 (80%) | 20 | 60.25 | 60 | 2 | -2.4 | -2.0 | -2.0 | |
| *IRF7* | IRF7 F | CGTGATGTTGCAAGACAACTCA | 3 | 22 | 57.22 | 45.45 | 1 | -2.4 | -5.7 | -2.4 | 96 |
| | IRF7 R | TGGTTAACGCCTGGGTCTCT | 3 (25%) / 4 (75%) | 20 | 57.72 | 55 | 1 | -2.4 | -4.3 | 0.0 | |
| *NFKB1* | NFKB1 F | GGGACAGTGTCTTACACTTAGCAATC | 13 (26.9%) /14 (73.1%) | 26 | 59.8 | 46.15 | 1 | -1.3 | -4.0 | -4.0 | 90 |
| | NFKB1 R | CATCAGAAATCAAGCCAGATGTG | 14 | 23 | 55.79 | 43.48 | 1 | -1.3 | -2.1 | -2.1 | |
| *RELA* | RELA F | AGAGTCCTTTCAATGGCCCCACCG | 7 (66.7%) / 8 (33.3%) | 24 | 64.61 | 58.33 | 3 | -2.4 | -4.4 | 0.0 | 81 |
| | RELA R | GGATGGAAGTTGAGCTGCGGGA | 8 | 22 | 62.27 | 59.09 | 3 | -2.4 | -3.0 | 0.0 | |
| *IKBKB* | IKBKB F | TAATGAACGAAGACGAGAAGATGGT | 18 | 25 | 58.16 | 40 | 2 | -4.4 | 0.0 | 0.0 | 91 |
| | IKBKB R | ACCTTGCTACACGCAATCTTCAG | 18 (78.3%) /19 (21.7%) | 23 | 59.11 | 47.83 | 2 | -4.4 | -3.1 | -3.1 | |
| *CXCL10* | CXCL10 F | CGTGTTGAGATTATTGCCACAATG | 2 (54%) /3 (46%) | 24 | 57.13 | 41.67 | 1 | -2.3 | -1.7 | -1.7 | 184 |
| | CXCL10 R | GAGGTAGCTTCTCTCTGGTCCT | 4 | 22 | 57.76 | 54.55 | 1 | -2.3 | -3.0 | -1.3 | |
| *MX1* | MX1 F | GAAGATGGTTTATTCTGACTCG | 2 | 22 | 52.21 | 40.91 | 2 | -0.7 | -0.7 | -0.7 | 146 |
| | MX1 R | TTCTCCTCGTACTGGCTGT | 3 | 19 | 54.29 | 52.63 | 2 | -0.7 | -2.0 | 0.0 | |
| *OAS1* | OAS1 F | TGAAGAAGCAGCTCGGGAAAC | 8 | 21 | 58.11 | 52.38 | 1 | -1.8 | -3.0 | 0.0 | 198 |
| | OAS1 R | AGTAACTGTCTTTTCTGGGCAGC | 9 (22%) /10 (78%) | 23 | 58.72 | 47.83 | 1 | -1.8 | -1.1 | -1.1 | |
| *ISG15* | ISG15 F M | CACAGCCATGGGTGGAATC | 1 (47.4%) / 2 (52.6%) | 19 | 55.19 | 57.89 | 1 | -4.2 | -6.1 | -1.5 | 91 |

| | | | | | | | | | | | |
|---|---|---|---|---|---|---|---|---|---|---|---|
| | ISG15 R M | CAGCTCCGATAACAGCATGGA | 2 | 21 | 57.43 | 52.38 | 1 | -4.2 | -3 | -1.8 | |
| *IL-10* | IL-10 F | CTGCTGGAGGACTTTAAGGGT | 2 (85%) /3 (15%) | 21 | 56.54 | 52.38 | 3 | -3.5 | -0.8 | 0.0 | 187 |
| | IL-10 R | AGGGGAGAAATCGATGACAGC | 3 (24%) /4 (76%) | 21 | 57.06 | 52.38 | 3 | -3.5 | -5.0 | 0.0 | |
| *IL-1β* | IL1-β F | AGGATATGAGCCGAGAAGTGGT | 5 (82%) /6 (18%) | 22 | 58.08 | 50.00 | 2 | -1.8 | -0.5 | 0.0 | 125 |
| | IL1-β R | CCCTTTCATCACACAAGACAGGT | 6 | 23 | 58.39 | 47.83 | 2 | -1.8 | -1.3 | -1.3 | |
| *IL-6* | IL-6 F | TCTGGGTTCAATCAGGAGACCT | 3 (68%) /4 (32%) | 22 | 57.86 | 50.00 | 2 | -1.5 | -1.5 | -1.5 | 192 |
| | IL-6 R | AGGGGTGCTTACTTCTTCTGGT | 5 | 22 | 58.4 | 50.00 | 2 | -1.5 | 0.0 | 0.0 | |
| *IL-8* | IL-8 F | TGTGTGAAGCTGCAGTTCTGT | 1 (43%) /2 (57%) | 21 | 57.62 | 47.62 | 1 | -3.5 | -6.5 | -0.6 | 176 |
| | IL-8 R | GCAGACCTCTCTTCCATTGGC | 3 | 21 | 58.24 | 57.14 | 1 | -3.5 | -1.5 | -0.7 | |
| *IL-15* | IL-15 F | CAGCCTACAGAAGGTCATGAAGTAC TC | 2 (66.7%) / 3 (33.3%) | 27 | 61.09 | 48.15 | 1 | -2.4 | -4.9 | -1.3 | 93 |
| | IL-15 R | GGGTAACTCCTTAAGTATCGAAGAA GAG | 3 | 28 | 59.22 | 42.82 | 1 | -2.4 | -3.9 | -1.0 | |
| *IL-2* | IL-2 F* | AAACTCTCCAGGATGCTCAC* | 2 | 20 | 54.17 | 50 | 1 | -2.3 | -1.3 | 0.0 | 202 |
| | IL-2 R | TTTCAGATCCCTTCAGTTCC | 3 (50%) /4 (50%) | 20 | 51.6 | 45 | 1 | -2.3 | -2.0 | 0.0 | |
| *IL-4* | IL-4 F | CCCTGGTCTGCTTACTGGTTT | 1 | 21 | 57.10 | 52.38 | 2 | -2.5 | 0.0 | 0.0 | 168 |
| | IL-4 R | TCTCAGTCGTGTTCTTTGGGG | 2 (38%) /3 (62%) | 21 | 57.14 | 52.38 | 2 | -2.5 | 0.0 | 0.0 | |
| *IL-12p35* | IL-12p35 F | AATCACCTGGACCACCTCAGT | 2 | 21 | 57.88 | 52.38 | 1 | -1.5 | -1.5 | -1.1 | 140 |
| | IL-12p35 R | TCTAGGGTTTGTCTGGCCTTC | 2 (15%) /3 (85%) | 21 | 56.55 | 52.38 | 1 | -1.5 | -4.4 | -1.3 | |
| *TNF-α* | TNF-α F | TGGCCCAGACCCTCAGATCA | 2 (75%) /3 (25%) | 20 | 59.30 | 60.00 | 1 | -2.4 | -4.4 | 0.0 | 143 |
| | TNF-α R | TTCCAGCTTCACACCATTGGC | 4 | 21 | 58.63 | 52.38 | 1 | -2.4 | -3.0 | -1.5 | |
| *CCL2* | CCL2 F | CCAGTAAGAAGATCCCCATGCA | 2 | 22 | 57.23 | 50 | 2 | -2 | -3.4 | 0.0 | 93 |
| | CCL2 R | GTGTGGTCTTGAAGATCACAGCTT | 2 (41.6%) / 3 (58.4%) | 24 | 59.15 | 45.83 | 2 | -2 | -3.0 | -2.7 | |
| *CCL3* | CCL3 F | GCTCAGCGTCATGCAGGTGCC | 1 | 21 | 63.98 | 66.67 | 3 | -2.4 | -3.4 | 0.0 | 113 |
| | CCL3 R | AGCAGGCGGTTGGGGTGTCAG | 2 | 21 | 64.45 | 66.67 | 3 | -2.4 | 0.0 | 0.0 | |
| *CXCL1* | CXCL1 F | CGTGCAGGGAATTCACTTCAA | 3 | 21 | 56.37 | 47.62 | 1 | -2.5 | -4.3 | -1.3 | 91 |
| | CXCL1 R | GAGAGTGGCTACGACTTCCGTTT | 3 (56.5%) / 4 (43.5%) | 23 | 60.15 | 52.17 | 1 | -2.5 | -1.9 | -1.9 | |
| *MIF* | MIF F | GCGAGTTGGTCGGTTCCTGTGTT | 1 | 23 | 63.19 | 56.52 | 1 | -2.9 | -1.8 | -1.8 | 176 |

| | | | | | | | | | | | |
|---|---|---|---|---|---|---|---|---|---|---|---|
| | MIF R | ACCACGTGCACTGCGATGTACT | 1 (9%) / 2 (91%) | 22 | 62.18 | 54.55 | 1 | -2.9 | -6.8 | 0.0 | |
| *CASP1* | CASP1 F | ACTCCACCAAGACCTCAACCAGT | 2 | 23 | 60.97 | 52.17 | 1 | -1.5 | -0.8 | -0.8 | 164 |
| | CASP1 R | GGGTAAATCTCCGCTGACTTCTCG | 3 (62.5%) / 4 (37.5%) | 24 | 60.92 | 54.17 | 1 | -1.5 | 0.0 | 0.0 | |
| *CASP10* | CASP10 F | CGGTAGCCACGGGAACTGAGTCAT | 5 | 24 | 64.12 | 58.33 | 1 | -2.4 | -0.9 | 0 | 107 |
| | CASP10 R | ATCTTGCCAGGACCCCTCCGAT | 5 (18.2%) / 6 (82.8%) | 22 | 62.6 | 59.09 | 1 | -2.4 | -1.3 | -1.3 | |
| *CYLD* | CYLD F | TCGGGATGGTGGTCAGAATGGC | 17 (41%) / 18 (59%) | 22 | 62.02 | 59.09 | 3 | -2.4 | 0.0 | 0.0 | 135 |
| | CYLD R | AGTCTTCGTGCACAGCCCTGGAT | 18 | 23 | 63.83 | 56.52 | 3 | -2.4 | -6.8 | 0.0 | |
| *AZI2* | AZI2 F | TGAGCGTCTCCAGCGCTAA | 6 (68.4%) / 7 (31.6%) | 19 | 58.03 | 57.89 | 2 | -2.9 | -7.7 | -4.1 | 86 |
| | AZI2 R | CTGCACTTGCGTCACCAGAT | 6 | 20 | 57.95 | 55 | 2 | -2.9 | -3.4 | -1.1 | |
| *PACT* | PACT F | TGCAGTTCCTGACCCCTTAATG | 3 | 22 | 57.71 | 50 | 1 | -1.1 | -3.4 | -1.1 | 92 |
| | PACT R | GATGAATAGCCAGTTCCTGTAGTGA A | 3 (38.5%) / 4 (61.5%) | 26 | 58.84 | 42.31 | 1 | -1.1 | -1.1 | -1.1 | |
| *TBK1* | TBK1 F | GTACAGAAAGCAGAAAATGGACCA A | 7 | 25 | 58.05 | 40 | 2 | -0.7 | -2.0 | 0.0 | 81 |
| | TBK1 R | AACTTGAAGGCCCCGAGAAA | 7 (35%) / 8 (65%) | 20 | 56.42 | 50 | 2 | -0.7 | -4.4 | 0.0 | |
| *TRIM25* | TRIM25 F | GCCCGAGCTCCTACAGTATGC | 7 (81.0%) / 8 (19.0%) | 21 | 59.86 | 61.9 | 2 | -5.2 | -6.2 | 0.0 | 93 |
| | TRIM25 R | GAAGCGACGGTGTAGGTCTTG | 8 | 21 | 58.29 | 57.14 | 2 | -5.2 | -1.1 | -1.1 | |
| *NFKBIA* | NFKBIA F | TCCCTCTTTTCCCCGCAGGTT | 2 | 21 | 60.88 | 57.14 | 2 | -3.5 | -1.3 | -1.3 | 138 |
| | NFKBIA R | TGGAGTGGAGTCTGCTGCAGGT | 2 (40.1%) / 3 (59.9%) | 22 | 62.96 | 59.09 | 2 | -3.5 | -6.5 | -1.1 | |
| *TRADD* | TRADD F | CGGCCAGGAAGCAAGATG | 1 (38.9%) / 2 (61.1%) | 18 | 54.92 | 61.11 | 1 | -2.0 | -4.4 | 0.0 | 81 |
| | TRADD R | TGAAGACTCCACAAACAGGTATGC | 2 | 24 | 58.91 | 45.83 | 1 | -2.0 | 0.0 | 0.0 | |
| *CARD9* | CARD9 F | GGCAGTGCAAGGTCCTGAAC | 1 | 20 | 58.5 | 60 | 1 | -4.3 | -3.4 | -1.1 | 92 |
| | CARD9 R | CAGGAGCACACCCACTTTCC | 1 (45%) / 2 (55%) | 20 | 57.85 | 60 | 1 | -4.3 | -1.3 | -1.3 | |
| *PYCARD* | PYCARD F | CAAGCCAGCACCGCACTT | 2 (44.4%) / 3 (65.6%) | 18 | 57.69 | 61.11 | 1 | -1.1 | -0.5 | -0.5 | 105 |
| | PYCARD R | TCTGTCAGGACCTTCCCATACA | 3 | 22 | 57.69 | 50 | 1 | -1.1 | -1.3 | -1.3 | |
| *IFNLR1* | IFNLR1 F | CAGGGTGTGTGATCTGGAAGAG | 6 | 22 | 57.81 | 54.55 | 1 | -5.6 | -2.0 | -1.1 | 90 |
| | IFNLR1 R | GTCTGTGTCCAGAGAAATCCAGG | 6 (17.4%) / 7 (82.6%) | 23 | 58.29 | 52.17 | 1 | -5.6 | -2.4 | -2.4 | |

| | | | | | | | | | | | |
|---|---|---|---|---|---|---|---|---|---|---|---|
| *IFNAR1* | IFNAR1 F | TGCGAGGAAACCAAACCAGGAAAT | 9 (84.3%) / 10 (16.7%) | 23 | 61.12 | 45.83 | 1 | -2.9 | 0.0 | 0.0 | 83 |
| | IFNAR1 R | ACGACGACGATACAAAACACCGC | 11 | 24 | 61.75 | 52.17 | 1 | -2.9 | 0.0 | 0.0 | |

(a) For primers designed at exon-exon boundaries, the percentage of nucleotides annealing each exon respect to the total number of nucleotides of the primer is indicated.

The asterisk (*) indicate a primer described by Odbileg et al. (2008). Tm was calculated for each primer at 3 mM of free $Mg^{2+}$ concentration. bp, base pairs; C, cytosine; G, guanine; GC clamp, presence of a guanine or cytosine base in the last 5 bases (3′ end) of a primer; Tm, melting temperature.
