## Supplementary material for "High-throughput quantification of camelid cytokine mRNA expression in PBMCs by microfluidic qPCR technology": Table S3

**Table S3**. Performance of the primer pairs designed for the quantification of mRNA expression of camelid immune and reference genes.

| Gene | Species | Stimulus[(a)] | N. of standard dilutions | Slope | $R^2$ | Efficacy | Efficacy (%) |
|---|---|---|---|---|---|---|---|
| *GAPDH* | Alpaca | PHA/PMA-ionomycin | 5 | -3.7633 | 0.9991 | 1.843844242 | 84.38442419 |
| | Alpaca | Poly I:C | 5 | -3.7776 | 0.9987 | 1.839578562 | 83.95785616 |
| | Llama | PHA/PMA-ionomycin | 5 | -3.7589 | 0.9987 | 1.84516529 | 84.51652897 |
| | Dromedary | PHA/PMA-ionomycin | 5 | -3.7269 | 0.9941 | 1.854895787 | 85.48957873 |
| *HPRT1* | Alpaca | PHA/PMA-ionomycin | 5 | -3.7653 | 0.999 | 1.843245098 | 84.32450983 |
| | Alpaca | Poly I:C | 5 | -3.8365 | 0.998 | 1.82244421 | 82.24442096 |
| | Llama | PHA/PMA-ionomycin | 5 | -3.8478 | 0.998 | 1.819234853 | 81.92348528 |
| | Dromedary | PHA/PMA-ionomycin | 5 | -3.8954 | 0.9963 | 1.805980456 | 80.5980456 |
| *UbC* | Alpaca | PHA/PMA-ionomycin | 5 | -3.6832 | 0.9984 | 1.86854276 | 86.85427603 |
| | Alpaca | Poly I:C | 5 | -3.6604 | 0.999 | 1.875833063 | 87.58330633 |
| | Llama | PHA/PMA-ionomycin | 5 | -3.7495 | 0.9992 | 1.848001107 | 84.80011067 |
| | Dromedary | PHA/PMA-ionomycin | 5 | -3.7847 | 0.9985 | 1.837476252 | 83.74762521 |
| *IFN-α* | Alpaca | PHA/PMA-ionomycin | 5 | -3.6903 | 0.9987 | 1.866296661 | 86.62966611 |
| | Alpaca | Poly I:C | 5 | -3.6625 | 0.9962 | 1.875156599 | 87.51565993 |
| | Llama | PHA/PMA-ionomycin | 4 | -3.9079 | 0.9849 | 1.802569055 | 80.25690553 |
| | Dromedary | PHA/PMA-ionomycin | 5 | -3.4502 | 0.9812 | 1.94911862 | 94.91186196 |
| *IFN-β* | Alpaca | PHA/PMA-ionomycin | 4 | -3.9686 | 0.9965 | 1.786397192 | 78.63971925 |
| | Alpaca | Poly I:C | | | | Poorly expressed in these cells | |
| | Llama | PHA/PMA-ionomycin | | | | Poorly expressed in these cells | |
| | Dromedary | PHA/PMA-ionomycin | 4 | -3.6585 | 0.9453 | 1.876445982 | 87.64459819 |
| *IFN-γ* | Alpaca | PHA/PMA-ionomycin | 4 | -4.2675 | 0.9929 | 1.715257127 | 71.52571272 |
| | Alpaca | Poly I:C | 4 | -4.1922 | 0.9934 | 1.731961515 | 73.19615155 |
| | Llama | PHA/PMA-ionomycin | 5 | -3.9601 | 0.9975 | 1.788623267 | 78.86232665 |
| | Dromedary | PHA/PMA-ionomycin | 5 | -3.8653 | 0.9982 | 1.81431266 | 81.431266 |
| *IFN-λ1* | Alpaca | PHA/PMA-ionomycin | 4 | -3.4396 | 0.9674 | 1.953131482 | 95.31314822 |
| | Alpaca | Poly I:C | 3 | -3.4477 | 0.991 | 1.950062084 | 95.00620836 |
| | Llama | PHA/PMA-ionomycin | | | | Poorly expressed in these cells | |
| | Dromedary | PHA/PMA-ionomycin | | | | Poorly expressed in these cells | |

| | | | | | | | |
|---|---|---|---|---|---|---|---|
| *IFN-λ3* | Alpaca | PHA/PMA-ionomycin | | | | Not expressed in these cells | |
| | Alpaca | PHA/PMA-ionomycin | | | | Not expressed in these cells | |
| | Alpaca | Poly I:C | | | | Not expressed in these cells | |
| | Llama | PHA/PMA-ionomycin | | | | Not expressed in these cells | |
| | Dromedary | PHA/PMA-ionomycin | | | | Not expressed in these cells | |
| *RIG-1* | Alpaca | PHA/PMA-ionomycin | 5 | -3.7589 | 0.9994 | 1.84516529 | 84.51652897 |
| | Alpaca | Poly I:C | 5 | -3.9091 | 0.9972 | 1.802243046 | 80.22430465 |
| | Llama | PHA/PMA-ionomycin | 5 | -3.948 | 0.9978 | 1.791813509 | 79.18135088 |
| | Dromedary | PHA/PMA-ionomycin | 5 | -3.9758 | 0.9972 | 1.784521178 | 78.45211779 |
| *MDA5* | Alpaca | PHA/PMA-ionomycin | 5 | -3.831 | 0.9987 | 1.824015196 | 82.40151958 |
| | Alpaca | Poly I:C | 5 | -3.8674 | 0.9963 | 1.813725881 | 81.37258809 |
| | Llama | PHA/PMA-ionomycin | 5 | -3.9904 | 0.9938 | 1.78074381 | 78.07438104 |
| | Dromedary | PHA/PMA-ionomycin | 3 | -3.9036 | 0.886 | 1.803739385 | 80.3739385 |
| *MAVS* | Alpaca | PHA/PMA-ionomycin | 5 | -3.7212 | 0.9991 | 1.856652031 | 85.66520312 |
| | Alpaca | Poly I:C | 5 | -3.7162 | 0.9967 | 1.858198407 | 85.81984065 |
| | Llama | PHA/PMA-ionomycin | 5 | -3.7478 | 0.9932 | 1.848515953 | 84.85159529 |
| | Dromedary | PHA/PMA-ionomycin | 5 | -3.8378 | 0.9942 | 1.822073741 | 82.20737408 |
| *TLR3* | Alpaca | PHA/PMA-ionomycin | 4 | -4.048 | 0.9899 | 1.766182483 | 76.6182483 |
| | Alpaca | Poly I:C | 4 | -4.1274 | 0.992 | 1.746961273 | 74.69612732 |
| | Llama | PHA/PMA-ionomycin | 4 | -3.7874 | 0.9884 | 1.83667948 | 83.66794795 |
| | Dromedary | PHA/PMA-ionomycin | 4 | -4.149 | 0.9849 | 1.741894841 | 74.18948414 |
| *TLR7* | Alpaca | PHA/PMA-ionomycin | 5 | -3.8778 | 0.999 | 1.810832074 | 81.08320744 |
| | Alpaca | Poly I:C | 5 | -3.9386 | 0.9983 | 1.794309365 | 79.43093653 |
| | Llama | PHA/PMA-ionomycin | 5 | -3.7304 | 0.989 | 1.853820872 | 85.3820872 |
| | Dromedary | PHA/PMA-ionomycin | 5 | -3.7653 | 0.9903 | 1.843245098 | 84.32450983 |
| *NLRP3* | Alpaca | PHA/PMA-ionomycin | 5 | -3.7865 | 0.996 | 1.836944906 | 83.69449057 |
| | Alpaca | Poly I:C | 5 | -3.8443 | 0.9977 | 1.82022628 | 82.02262805 |
| | Llama | PHA/PMA-ionomycin | 5 | -3.8271 | 0.9938 | 1.825132727 | 82.51327274 |
| | Dromedary | PHA/PMA-ionomycin | 5 | -3.9738 | 0.9952 | 1.785041414 | 78.50414136 |
| *STAT1* | Alpaca | PHA/PMA-ionomycin | 5 | -3.9421 | 0.9992 | 1.79337826 | 79.337826 |
| | Alpaca | Poly I:C | 5 | -3.9938 | 0.9975 | 1.779869256 | 77.98692563 |
| | Alpaca | Poly I:C | 4 | -3.9284 | 0.999 | 1.797035111 | 79.70351106 |

| | | | | | | | |
|---|---|---|---|---|---|---|---|
| | Llama | PHA/PMA-ionomycin | 4 | -4.0362 | 0.9954 | 1.769122036 | 76.91220357 |
| | Dromedary | PHA/PMA-ionomycin | 5 | -3.8908 | 0.999 | 1.807243002 | 80.72430025 |
| *IRF3* | Alpaca | PHA/PMA-ionomycin | 5 | -3.826 | 0.9969 | 1.825448464 | 82.54484645 |
| | Alpaca | Poly I:C | 5 | -3.9064 | 0.9956 | 1.802976931 | 80.2976931 |
| | Llama | PHA/PMA-ionomycin | 5 | -3.8687 | 0.9955 | 1.813363051 | 81.3363051 |
| | Dromedary | PHA/PMA-ionomycin | 5 | -3.8588 | 0.9933 | 1.816134137 | 81.61341366 |
| *IRF5* | Alpaca | PHA/PMA-ionomycin | 5 | -3.831 | 0.9991 | 1.824015196 | 82.40151958 |
| | Alpaca | Poly I:C | 5 | -3.8493 | 0.998 | 1.818810672 | 81.88106722 |
| | Llama | PHA/PMA-ionomycin | 5 | -3.8443 | 0.9992 | 1.82022628 | 82.02262805 |
| | Dromedary | PHA/PMA-ionomycin | 5 | -3.8215 | 0.9955 | 1.826742579 | 82.67425794 |
| *IRF7* | Alpaca | PHA/PMA-ionomycin | 5 | -3.7724 | 0.9982 | 1.841124834 | 84.11248345 |
| | Alpaca | Poly I:C | 5 | -3.7783 | 0.9972 | 1.839370834 | 83.93708336 |
| | Llama | PHA/PMA-ionomycin | 5 | -3.7682 | 0.9979 | 1.842377815 | 84.23778149 |
| | Dromedary | PHA/PMA-ionomycin | 3 | -3.84 | 0.9621 | 1.821447536 | 82.14475364 |
| *NFKB1* | Alpaca | PHA/PMA-ionomycin | 5 | -3.7739 | 0.9981 | 1.840678224 | 84.06782235 |
| | Alpaca | Poly I:C | 5 | -3.8145 | 0.9993 | 1.828763549 | 82.87635487 |
| | Llama | PHA/PMA-ionomycin | 5 | -3.7738 | 0.9979 | 1.840707983 | 84.07079832 |
| | Dromedary | PHA/PMA-ionomycin | 5 | -3.7852 | 0.9979 | 1.83732859 | 83.73285898 |
| *RELA* | Alpaca | PHA/PMA-ionomycin | 5 | -3.9117 | 0.9989 | 1.801537582 | 80.15375821 |
| | Alpaca | Poly I:C | 5 | -3.9084 | 0.998 | 1.802433187 | 80.24331868 |
| | Llama | PHA/PMA-ionomycin | 5 | -3.9647 | 0.9981 | 1.787417038 | 78.74170385 |
| | Dromedary | PHA/PMA-ionomycin | 5 | -3.9185 | 0.9966 | 1.799698248 | 79.96982475 |
| *IKBKB* | Alpaca | PHA/PMA-ionomycin | 5 | -3.7803 | 0.9994 | 1.838777878 | 83.87778779 |
| | Alpaca | Poly I:C | 5 | -3.8002 | 0.997 | 1.832922258 | 83.29222579 |
| | Llama | PHA/PMA-ionomycin | 5 | -3.8024 | 0.9984 | 1.832279804 | 83.22798039 |
| | Dromedary | PHA/PMA-ionomycin | 5 | -3.806 | 0.9961 | 1.831230602 | 83.12306021 |
| *CXCL10* | Alpaca | PHA/PMA-ionomycin | 5 | -3.8401 | 0.9982 | 1.821419095 | 82.14190947 |
| | Alpaca | Poly I:C | 5 | -4.0118 | 0.9865 | 1.775271043 | 77.5271043 |
| | Llama | PHA/PMA-ionomycin | 5 | -3.8842 | 0.9994 | 1.809061254 | 80.90612544 |
| | Dromedary | PHA/PMA-ionomycin | 5 | -3.854 | 0.9944 | 1.81748435 | 81.74843495 |
| *MX1* | Alpaca | PHA/PMA-ionomycin | 5 | -3.803 | 0.9984 | 1.832104757 | 83.21047572 |
| | Alpaca | Poly I:C | 5 | -3.856 | 0.9983 | 1.81692123 | 81.69212303 |

| | | | | | | | |
|---|---|---|---|---|---|---|---|
| | Llama | PHA/PMA-ionomycin | 5 | -3.7916 | 0.9977 | 1.835442996 | 83.54429957 |
| | Dromedary | PHA/PMA-ionomycin | 5 | -3.8606 | 0.9988 | 1.81562893 | 81.56289305 |
| *OAS1* | Alpaca | PHA/PMA-ionomycin | 5 | -3.7314 | 0.9993 | 1.853514238 | 85.3514238 |
| | Alpaca | Poly I:C | 5 | -3.8802 | 0.9929 | 1.810167129 | 81.01671293 |
| | Llama | PHA/PMA-ionomycin | 5 | -3.6702 | 0.9851 | 1.872684931 | 87.26849309 |
| | Dromedary | PHA/PMA-ionomycin | 5 | -3.8741 | 0.9978 | 1.811859292 | 81.18592922 |
| *ISG15* | Alpaca | PHA/PMA-ionomycin | 5 | -3.7632 | 0.9992 | 1.843874221 | 84.38742209 |
| | Alpaca | Poly I:C | 5 | -3.862 | 0.9938 | 1.815236415 | 81.5236415 |
| | Llama | PHA/PMA-ionomycin | 5 | -3.8145 | 0.9961 | 1.828763549 | 82.87635487 |
| | Dromedary | PHA/PMA-ionomycin | 5 | -3.813 | 0.9957 | 1.829197871 | 82.91978705 |
| *IL-10* | Alpaca | PHA/PMA-ionomycin | 5 | -3.8501 | 0.9923 | 1.818584618 | 81.8584618 |
| | Alpaca | Poly I:C | 5 | -3.9612 | 0.9951 | 1.788334492 | 78.83344918 |
| | Llama | PHA/PMA-ionomycin | 5 | -3.7541 | 0.9918 | 1.846611048 | 84.66110479 |
| | Dromedary | PHA/PMA-ionomycin | 5 | -3.7373 | 0.9865 | 1.851709466 | 85.1709466 |
| *IL-1β* | Alpaca | PHA/PMA-ionomycin | 5 | -3.7876 | 0.9958 | 1.836620518 | 83.66205182 |
| | Alpaca | Poly I:C | 5 | -3.8174 | 0.9987 | 1.82792512 | 82.79251195 |
| | Llama | PHA/PMA-ionomycin | 5 | -3.8103 | 0.9993 | 1.829980772 | 82.9980772 |
| | Dromedary | PHA/PMA-ionomycin | 5 | -3.8102 | 0.9915 | 1.830009796 | 83.00097961 |
| *IL-6* | Alpaca | PHA/PMA-ionomycin | 5 | -3.7502 | 0.9819 | 1.847789288 | 84.77892885 |
| | Alpaca | Poly I:C | 5 | -3.8015 | 0.9893 | 1.832542509 | 83.25425089 |
| | Llama | PHA/PMA-ionomycin | 5 | -3.908 | 0.9973 | 1.802541878 | 80.2541878 |
| | Dromedary | PHA/PMA-ionomycin | 5 | -3.8766 | 0.9974 | 1.811164947 | 81.11649475 |
| *IL-8* | Alpaca | PHA/PMA-ionomycin | 5 | -3.8137 | 0.9981 | 1.828995132 | 82.89951317 |
| | Alpaca | Poly I:C | 5 | -3.6108 | 0.997 | 1.892112422 | 89.21124222 |
| | Llama | PHA/PMA-ionomycin | 5 | -3.5948 | 0.9969 | 1.897490424 | 89.74904236 |
| | Dromedary | PHA/PMA-ionomycin | 5 | -3.6951 | 0.9972 | 1.864784585 | 86.47845846 |
| *IL-15* | Alpaca | PHA/PMA-ionomycin | 5 | -3.7972 | 0.9971 | 1.833799895 | 83.3799895 |
| | Alpaca | Poly I:C | 5 | -3.7939 | 0.9985 | 1.834767386 | 83.47673857 |
| | Llama | PHA/PMA-ionomycin | 5 | -4.0218 | 0.9984 | 1.772739352 | 77.27393517 |
| | Dromedary | PHA/PMA-ionomycin | 5 | -3.8056 | 0.9962 | 1.831347052 | 83.13470524 |
| *IL-2* | Alpaca | PHA/PMA-ionomycin | 5 | -3.5408 | 0.9635 | 1.916117121 | 91.61171214 |
| | Alpaca | Poly I:C | 4 | -3.9649 | 0.9809 | 1.787364676 | 78.73646757 |

| | | | | | | | |
|---|---|---|---|---|---|---|---|
| | Llama | PHA/PMA-ionomycin | 5 | -4.0276 | 0.9982 | 1.771278377 | 77.12783768 |
| | Dromedary | PHA/PMA-ionomycin | 5 | -3.9098 | 0.9992 | 1.802052994 | 80.20529943 |
| *IL-4* | Alpaca | PHA/PMA-ionomycin | 3 | -3.443 | 0.9889 | 1.951840746 | 95.18407458 |
| | Alpaca | Poly I:C | 5 | -3.3234 | 0.9672 | 1.999386116 | 99.93861165 |
| | Llama | PHA/PMA-ionomycin | 5 | -3.9908 | 0.9972 | 1.780640822 | 78.0640822 |
| | Dromedary | PHA/PMA-ionomycin | 5 | -3.9975 | 0.992 | 1.778919713 | 77.89197129 |
| *IL-12p35* | Alpaca | PHA/PMA-ionomycin | 4 | -3.9231 | 0.982 | 1.798458668 | 79.84586681 |
| | Alpaca | Poly I:C | 3 | -3.2992 | 0.9833 | 2.00957298 | 100.957298 |
| | Llama | PHA/PMA-ionomycin | 4 | -3.7959 | 0.966 | 1.834180766 | 83.4180766 |
| | Dromedary | PHA/PMA-ionomycin | 5 | -3.3967 | 0.973 | 1.969714997 | 96.97149969 |
| *TNF-α* | Alpaca | PHA/PMA-ionomycin | 5 | -3.6676 | 0.9945 | 1.873517995 | 87.35179954 |
| | Alpaca | Poly I:C | 5 | -3.8683 | 0.9925 | 1.813474657 | 81.34746573 |
| | Llama | PHA/PMA-ionomycin | 5 | -3.8436 | 0.9983 | 1.820424848 | 82.04248476 |
| | Dromedary | PHA/PMA-ionomycin | 5 | -3.9524 | 0.9945 | 1.790650502 | 79.06505022 |
| *CCL2* | Alpaca | PHA/PMA-ionomycin | 5 | -3.7174 | 0.9972 | 1.85782678 | 85.78267795 |
| | Alpaca | Poly I:C | 5 | -3.6299 | 0.9976 | 1.885774171 | 88.57741706 |
| | Llama | PHA/PMA-ionomycin | 5 | -3.8695 | 0.9982 | 1.813139928 | 81.31399283 |
| | Dromedary | PHA/PMA-ionomycin | 5 | -3.7671 | 0.9975 | 1.842706579 | 84.27065792 |
| *CCL3* | Alpaca | PHA/PMA-ionomycin | 5 | -3.7709 | 0.9978 | 1.841571909 | 84.15719093 |
| | Alpaca | Poly I:C | 5 | -3.829 | 0.9989 | 1.824587919 | 82.4587919 |
| | Llama | PHA/PMA-ionomycin | 5 | -3.7693 | 0.9986 | 1.842049301 | 84.2049301 |
| | Dromedary | PHA/PMA-ionomycin | 5 | -3.81 | 0.9978 | 1.83006785 | 83.00678502 |
| *CXCL1* | Alpaca | PHA/PMA-ionomycin | 5 | -3.7946 | 0.9995 | 1.834561977 | 83.45619771 |
| | Alpaca | Poly I:C | 5 | -3.7309 | 0.9984 | 1.853667528 | 85.36675281 |
| | Llama | PHA/PMA-ionomycin | 5 | -3.7632 | 0.9986 | 1.843874221 | 84.38742209 |
| | Dromedary | PHA/PMA-ionomycin | 5 | -3.7908 | 0.9975 | 1.835678241 | 83.5678241 |
| *MIF* | Alpaca | PHA/PMA-ionomycin | 5 | -3.9039 | 0.997 | 1.803657626 | 80.36576258 |
| | Alpaca | Poly I:C | 5 | -3.8017 | 0.998 | 1.832484116 | 83.2484116 |
| | Llama | PHA/PMA-ionomycin | 5 | -3.9286 | 0.9985 | 1.796981489 | 79.69814887 |
| | Dromedary | PHA/PMA-ionomycin | 5 | -3.7621 | 0.989 | 1.844204127 | 84.42041274 |
| *CASP1* | Alpaca | PHA/PMA-ionomycin | 5 | -3.8193 | 0.9916 | 1.827376702 | 82.73767023 |
| | Alpaca | Poly I:C | 5 | -3.7936 | 0.9971 | 1.834855448 | 83.48554481 |

| | | | | | | | |
|---|---|---|---|---|---|---|---|
| | Llama | PHA/PMA-ionomycin | 5 | -3.8819 | 0.9937 | 1.809696771 | 80.96967714 |
| | Dromedary | PHA/PMA-ionomycin | 4 | -3.722 | 0.9964 | 1.856405116 | 85.64051159 |
| *CASP10* | Alpaca | PHA/PMA-ionomycin | 5 | -3.7891 | 0.9972 | 1.836178567 | 83.61785674 |
| | Alpaca | Poly I:C | 5 | -3.7671 | 0.9971 | 1.842706579 | 84.27065792 |
| | Llama | PHA/PMA-ionomycin | 5 | -3.8029 | 0.9967 | 1.832133927 | 83.21339266 |
| | Dromedary | PHA/PMA-ionomycin | 5 | -3.8375 | 0.9967 | 1.822159205 | 82.21592046 |
| *CYLD* | Alpaca | PHA/PMA-ionomycin | 5 | -3.8277 | 0.9986 | 1.824960607 | 82.49606066 |
| | Alpaca | Poly I:C | 5 | -3.8513 | 0.9982 | 1.818245766 | 81.82457656 |
| | Llama | PHA/PMA-ionomycin | 5 | -3.8544 | 0.9975 | 1.817371665 | 81.7371665 |
| | Dromedary | PHA/PMA-ionomycin | 5 | -3.8135 | 0.9909 | 1.829053047 | 82.90530472 |
| *AZI2* | Alpaca | PHA/PMA-ionomycin | 5 | -3.7329 | 0.9984 | 1.85305469 | 85.305469 |
| | Alpaca | Poly I:C | 5 | -3.6647 | 0.9946 | 1.874449015 | 87.44490155 |
| | Llama | PHA/PMA-ionomycin | 5 | -3.7789 | 0.9947 | 1.839192861 | 83.91928609 |
| | Dromedary | PHA/PMA-ionomycin | 5 | -3.8561 | 0.9924 | 1.816893094 | 81.68930943 |
| *PACT* | Alpaca | PHA/PMA-ionomycin | 5 | -3.8744 | 0.9958 | 1.811775909 | 81.17759094 |
| | Alpaca | Poly I:C | 5 | -3.7357 | 0.9898 | 1.852198159 | 85.21981587 |
| | Llama | PHA/PMA-ionomycin | 5 | -3.8179 | 0.9969 | 1.82778073 | 82.77807303 |
| | Dromedary | PHA/PMA-ionomycin | 5 | -3.9467 | 0.9941 | 1.792157765 | 79.21577654 |
| *TBK1* | Alpaca | PHA/PMA-ionomycin | 5 | -3.8119 | 0.9972 | 1.829516656 | 82.95166561 |
| | Alpaca | Poly I:C | 5 | -3.9139 | 0.9968 | 1.800941598 | 80.09415981 |
| | Llama | PHA/PMA-ionomycin | 5 | -3.7734 | 0.9964 | 1.840827042 | 84.08270424 |
| | Dromedary | PHA/PMA-ionomycin | 5 | -3.8847 | 0.9962 | 1.808923228 | 80.89232277 |
| *TRIM25* | Alpaca | PHA/PMA-ionomycin | 5 | -3.7586 | 0.9985 | 1.845255508 | 84.52555082 |
| | Alpaca | Poly I:C | 5 | -3.7857 | 0.9973 | 1.837180978 | 83.71809783 |
| | Llama | PHA/PMA-ionomycin | 5 | -3.7291 | 0.9991 | 1.854219818 | 85.4219818 |
| | Dromedary | PHA/PMA-ionomycin | 5 | -3.8824 | 0.9958 | 1.809558532 | 80.95585325 |
| *NFKBIA* | Alpaca | PHA/PMA-ionomycin | 5 | -3.8608 | 0.9952 | 1.815572834 | 81.55728342 |
| | Alpaca | Poly I:C | 5 | -3.8294 | 0.9974 | 1.824473312 | 82.44733121 |
| | Llama | PHA/PMA-ionomycin | 5 | -3.773 | 0.9976 | 1.840946134 | 84.09461345 |
| | Dromedary | PHA/PMA-ionomycin | 5 | -3.8332 | 0.9913 | 1.823386098 | 82.33860978 |
| *TRADD* | Alpaca | PHA/PMA-ionomycin | 5 | -3.7986 | 0.9951 | 1.833390106 | 83.33901062 |
| | Alpaca | Poly I:C | 5 | -3.888 | 0.996 | 1.808013405 | 80.80134049 |

| | | | | | | | |
|---|---|---|---|---|---|---|---|
| | Llama | PHA/PMA-ionomycin | 5 | -3.8988 | 0.9896 | 1.80504975 | 80.50497496 |
| | Dromedary | PHA/PMA-ionomycin | 5 | -3.8443 | 0.9952 | 1.82022628 | 82.02262805 |
| *CARD9* | Alpaca | PHA/PMA-ionomycin | 5 | -3.6531 | 0.9973 | 1.87819254 | 87.81925396 |
| | Alpaca | Poly I:C | 5 | -3.6925 | 0.9923 | 1.865602986 | 86.56029859 |
| | Llama | PHA/PMA-ionomycin | 5 | -3.8609 | 0.9792 | 1.815544789 | 81.55447889 |
| | Dromedary | PHA/PMA-ionomycin | 4 | -3.8076 | 0.9921 | 1.83076512 | 83.07651195 |
| *PYCARD* | Alpaca | PHA/PMA-ionomycin | 5 | -3.7062 | 0.9989 | 1.861307565 | 86.13075654 |
| | Alpaca | Poly I:C | 5 | -3.8105 | 0.9952 | 1.82992273 | 82.99227297 |
| | Llama | PHA/PMA-ionomycin | 5 | -3.7075 | 0.9697 | 1.860902132 | 86.09021317 |
| | Dromedary | PHA/PMA-ionomycin | 5 | -3.9313 | 0.9971 | 1.796258283 | 79.62582833 |
| *IFNLR1* | Alpaca | PHA/PMA-ionomycin | 5 | -3.7347 | 0.9983 | 1.85250387 | 85.25038699 |
| | Alpaca | Poly I:C | 5 | -3.8927 | 0.9945 | 1.806721047 | 80.67210472 |
| | Llama | PHA/PMA-ionomycin | 5 | -3.7049 | 0.9968 | 1.861713372 | 86.17133721 |
| | Dromedary | PHA/PMA-ionomycin | 3 | -3.9184 | 0.9293 | 1.799725237 | 79.97252367 |
| *IFNAR1* | Alpaca | PHA/PMA-ionomycin | 5 | -4.245 | 0.9958 | 1.720169569 | 72.01695685 |
| | Alpaca | Poly I:C | 5 | -4.3844 | 0.9953 | 1.690757721 | 69.07577213 |
| | Llama | PHA/PMA-ionomycin | 5 | -4.3109 | 0.9939 | 1.705965021 | 70.59650213 |
| | Dromedary | PHA/PMA-ionomycin | | | | Poorly expressed in these cells | |

(a) Each primer pair was validated using a mixture of cDNAs from alpaca, dromedary camel and llama PBMCs stimulated with PHA, PMA-ionomicyn or PolyI:C. PBMCs, peripheral blood mononuclear cells; PHA, phytohemagglutinin; PMA, phorbol 12-myristate 13-acetate.
